# Reliability of high-quantity human brain organoids for modeling microcephaly, glioma invasion, and drug screening

**DOI:** 10.1101/2023.10.29.564523

**Authors:** Anand Ramani, Giovanni Pasquini, Niklas J. Gerkau, Omkar Suhas Vinchure, Elke Gabriel, Ina Rothenaigner, Sean Lin, Nazlican Altinisk, Dhanasekaran Rathinam, Ali Mirsaidi, Olivier Goureau, Lucia Ricci-Vitiani Giorgio, Q. d’alessandris, Roberto Pallini, Bernd Wollnik, Alysson Muotri, Nathalie Jurisch-Yaksi, Christine R. Rose, Volker Busskamp, Kamyar Hadian, Jay Gopalakrishnan

## Abstract

Brain organoids offer unprecedented insights into brain development and disease modeling and hold promise for drug screening. Significant hindrances, however, are morphological and cellular heterogeneity, inter-organoid size differences, cellular stress, and poor reproducibility. Here, we describe a method that reproducibly generates thousands of organoids across multiple iPSC lines. These High Quantity brain organoids (Hi-Q brain organoids) exhibit reproducible cytoarchitecture, cell diversity, and functionality, are free from ectopically active cellular stress pathways, and allow cryopreservation and re-culturing. Using patient-derived Hi-Q brain organoids, we recapitulated distinct forms of microcephaly pathogenesis: primary microcephaly due to a mutation in centrosomal CDK5RAP2 and progeria-associated microcephaly in Cockayne syndrome. When modeling glioma invasion, hi-Q brain organoids displayed a similar invasion pattern for a given patient-derived glioma cell line. This enabled a medium-throughput screen to identify Selumetinib and Fulvestrant, which also perturbed glioma invasion in vivo. Thus, the Hi-Q approach can easily be adapted to reliably harness brain organoids’ utility for personalized disease modeling and drug discovery.

---

Recent developments in three-dimensional culturing of pluripotent stem cells and tissue engineering have led to the generation of 3D brain organoids, which offer unique opportunities to recapitulate various aspects of brain development and disease ^1,2^ ^3^ ^4^. In particular, brain organoids derived from healthy iPSCs have modeled early embryonic brain development ^5–7^. When cultured long-term, brain organoids generate more advanced cell types and functionally active neuronal networks ^3,8^. Besides recapitulating development, brain organoids hold value in modeling brain disorders when using patient-specific iPSCs or perturbing target genes by genome editing technologies ^2,9,10^. If generated on a large scale with reproducible quality, brain organoids can also offer drug screening assays to identify therapeutic compounds.

Several methods have been developed to culture brain organoids tailored to specific questions of interest. However, regardless of the protocol, significant challenges limit their application for reliable disease modeling and drug screening. These include morphological and regional heterogeneity, inter-organoid size differences, ectopically induced activation of cellular stress pathways, and, most importantly, the limited quantity of organoids per batch ^11,12^. Thus, it is essential to establish a scalable approach to generate whole-brain organoids of reproducible qualities and quantities capable of modeling disorders and screening drugs.

Most methods include the embryoid body (EB) as an intermediate stage before the neuroectoderm is further differentiated ^13,14^. However, EB generation typically requires manually embedding pluripotent cells in an extracellular matrix. This process can cause size differences between organoids and significantly limits the number of organoids one can generate per batch ^15^. As a result, organoids can often remarkably differ in size, morphology, and cellular diversity.

Here, we describe a simplified culturing method to induce direct differentiation of iPSCs into the neural epithelium, omitting the EB and using an extracellular matrix. Notably, by using custom-designed, coating-free, pre-patterned microwells, we have complete control over the sizes of early-stage neurospheres. Following a quick transfer to spinner-flask bioreactors, our method generates brain organoids in high quantities (referred to hereafter as Hi-Q brain organoids). This Hi-Q approach is highly efficient as we can culture several hundreds of brain organoids within a batch from multiple iPSC lines. Furthermore, Hi-Q brain organoids exhibit similar cell diversity, cytoarchitectures, maturation time, and functionality. Importantly, brain organoids generated with this platform displayed minimum to no ectopically activated cellular stress pathways, which has been previously shown to impair cell-type specification^13^. We report that Hi-Q brain organoids can also be successfully cryopreserved and re-cultured.

Human brain organoids offer the unique opportunity to underpin the pathomechanisms of human microcephaly. In particular, brain organoids derived from microcephaly patient iPSCs could recapitulate the early defects causing neocortex malformation ^9,16,17^. However, no standard method is optimized to model various microcephaly phenotypes robustly. Hi-Q brain organoids described here could recapitulate the phenotypes of both primary microcephaly due to a mutation in the centrosomal protein CDK5RAP2 and neurological phenotypes of Cockayne syndrome due to DNA damage. Finally, to assess the applicability of Hi-Q brain organoids for drug screening, first, we modeled glioma invasion by fusing patient-derived glioma stem cells (GSCs) to Hi-Q brain organoids. We ultimately used the GSC-invading Hi-Q brain organoids for a medium-throughput drug screening. Applying machine-learned algorithms and automated imaging, we identified Selumetinib and Fulvestrant as potent inhibitors, which could also impair GSC invasion in brain organoids and *in vivo* mouse xenografts.

## Results

### Generation of Hi-Q brain organoids

Conventionally, most unguided brain organoid differentiation methods included manual embedding of iPSCs in Matrigel. These embedded objects were then processed into the EB generation before differentiation into the neurospheres ^5,6,18,19^. As objects in culture dishes, EBs and neurospheres tend to exhibit high heterogeneity with variable shapes and sizes. We reasoned that differentiating iPSCs into neurospheres in a confined space could restrict heterogeneity and lead to the generation of uniform-sized brain organoids with an increased degree of homogeneity and, thus, cell diversity. To do this, we aimed to differentiate iPSCs into neurospheres in a confined space of a microwell equipped with a round bottom satisfying two prerequisites. First, microwells allow identical diffusion conditions for all spheres and unique physiological sphere formation. Second, microwell material does not require precoating that could dampen the cells’ attachment or a centrifugation step that forces cell pelleting. For this purpose, we fabricated a 24-well custom-designed spherical plate using a medical-grade, inert Cyclo-Olefin-Copolymer (COC), which offers an ideal surface property. The plate consists of 24 large wells, which are micropatterned to contain 185 equally sized microwells of 1×1mm at the opening and 180 µm in diameter at their round base **(Figure 1A).** Typically, each microwell exhibits an inverted pyramid shape with a rounded bottom. This geometry induces the seeded cells to form spheres through mutual adhesion. The advantage of this configuration is that it facilitates cell-cell communication. Importantly, our spherical plate has a geometry that allows precise control of sphere size, resulting in higher standardization and reproducibility combined with a high quantity of uniform-sized spheres.

**Figure 1:**
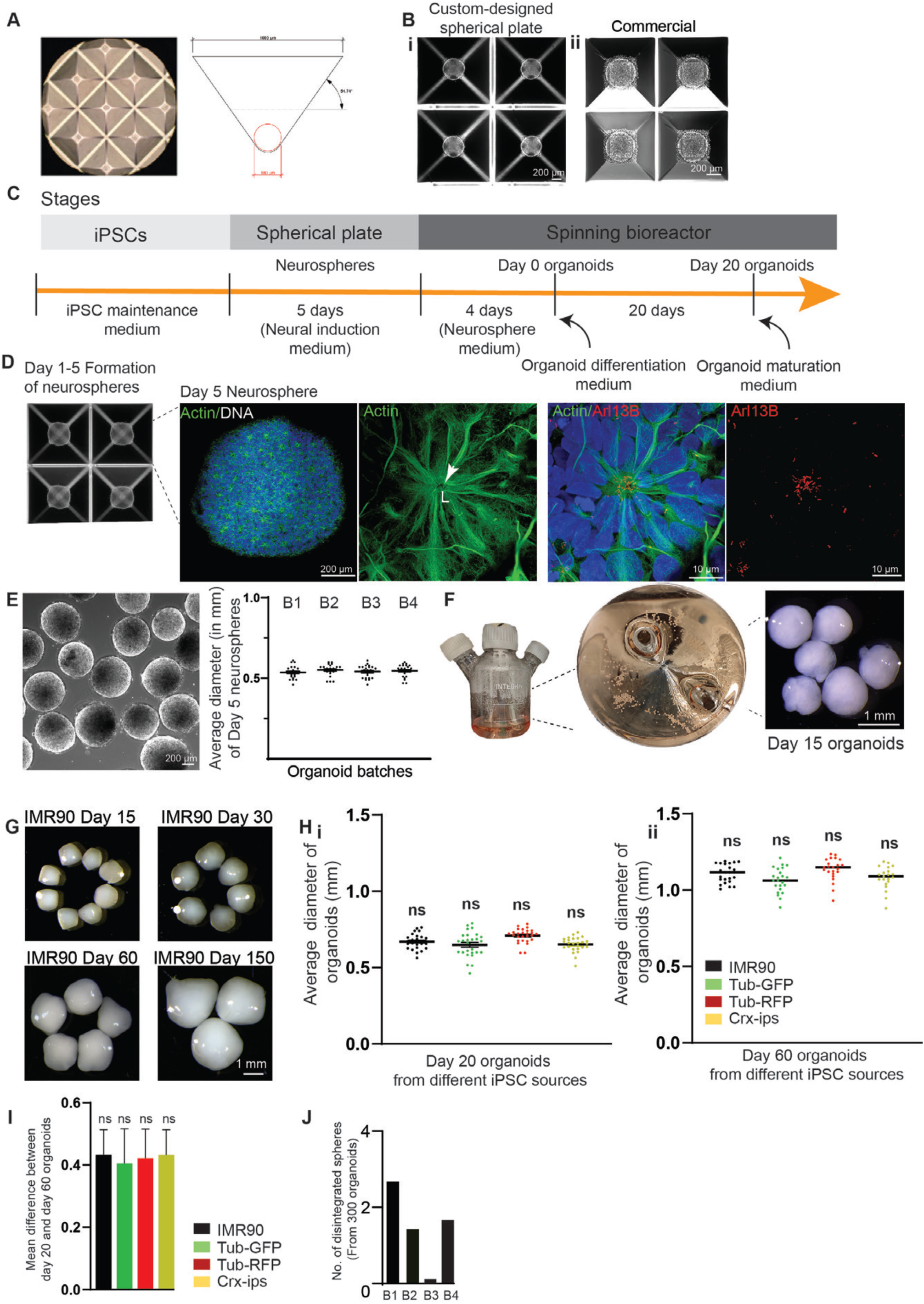
Generation of Hi-Q brain organoids. **A.** Microscopic view showing a custom-designed spherical plate with microwells and schematic view of the inverted pyramid-like microwell. The angles and measurements are shown. **B.** iPSCs settle and form spheres readily in the microwells of the spherical plate **(i)** but not efficiently in a commercially available plate (AggreWell 800) 24 hours after seeding **(ii).** Panels show the scale bar. **C.** Different stages of Hi-Q brain organoid generation. **D.** Formation of neurospheres in the microwells of the spherical plate. The magnified image at the right shows a representative neurosphere. Magnified panels show a neural rosette stained with actin (green) with primary cilia emanating into the lumen (L) at the apical side of the rosette marked by Arl13B (Red). Panels show the scale bar. **E.** A group of neurospheres. The graph at right shows no significant difference between the organoid diameter across four independent batches. At least twenty (n=20) randomly chosen neurospheres were measured from each batch. Statistical analysis was carried out by One-way ANOVA, followed by Tukey’s multiple comparisons test. Data presented as mean ± SEM. The cell line used is IMR90. The panel shows the scale bar. **F.** Maturation of neurospheres into Hi-Q brain organoids in spinner flasks. Macroscopic images show a group of organoids. The panel shows the scale bar. **G.** Hi-Q brain organoids increase in size progressively over time from day 15 to day 180. The panel shows the scale bar. **H.** The average diameter of twenty-day **(i)** and fifty-day **(ii)** old organoids were differentiated from four independent iPSC lines. Note that there is no significant difference among the different iPSCs donors within each time point. At least twenty-five (n=25) randomly chosen organoids were analyzed across three independent batches (N=3). Statistical analysis was carried out by One-way ANOVA, followed by Tukey’s multiple comparisons test. Data presented as mean ± SEM. **I.** The graph shows no size difference between twenty-day and fifty-day-old organoids, showing a regulated growth rate of the organoids between two time points. At least twenty-five (n=25) randomly chosen organoids were analyzed across three independent batches (N=3). Statistical analysis was carried out by One-way ANOVA, followed by Tukey’s multiple comparisons test. Data presented as mean ± SEM. **J.** The bar diagram quantifies the number of disintegrated organoids in each batch. At least three hundred (n=300) organoids were randomly sampled across four independent batches (N=4). One-way ANOVA carried out statistical analysis.

We noticed that iPSCs readily settled within a day of plating in our spherical plate, even without a centrifugation step. In contrast, the iPSCs did not settle well in commercially available microwell plates that required pre-coating. This finding suggests that our spherical plate may provide a more suitable environment for sphere formation **(Figure 1B).** Typically, using our spherical plates, we could differentiate 10,000 iPSCs into uniform-sized 3D neurospheres within each microwell. While using a Rho-kinase (ROCK) inhibitor at this stage for an extended period will alleviate cell death, prolonged exposure could change the cell’s metabolism and induce the meso-endodermal differentiation pathway^20,21^. Indeed, prolonged use of ROCK inhibitors is associated with generating of organoids with ectopically active cellular stress pathways ^13^. Therefore, after 24 hours of initial culturing, we omitted the ROCK inhibitor.

On day 5, we noticed uniform-sized neurospheres **(Figure 1C).** Imaging them revealed that they are highly similar from well to well, exhibiting characteristic neural rosette organization with primary cilia emanating apically into the lumen. Furthermore, neurosphere size remained consistent across several independent batches **(Figure 1D).** We then transferred these uniform-sized Matrigel-free neurospheres to spinner bioreactors containing 75 ml neurosphere medium **(Figure 1E, Method table 1)**. After culturing for four days, we switched to a brain organoid differentiation medium containing 5 µM SB431542 and 0.5 μM Dorsomorphin, inhibitors of TGF-β and BMP pathways to initiate an undirected neural differentiation. Twenty days later, we switched to a brain organoid maturation medium and cultured organoids until day 150 with a constant spinning rate of 25 RPM without noticing disintegrated spheroids **(Movie 1).**

**Table 1.** Media composition and incubation.

| **Sr. No** | **Media Name** | **Media Composition/ company)** | **No. of Days in respective media** |
| --- | --- | --- | --- |
| 1 | iPSC maintenance medium | mTeSR-1 (Stem Cell Technologies) | Variable (upto 80% confluency) |
| 2 | Neural Induction medium | STEM Diff Neural induction medium (NIM) (Stem cell technologies, Catalog # 05835) | 5 |
| 3 | Wash media | DMEM/ F12 | - |
| 4 | Neurosphere medium | 1:1 DMEM/F12 and Neural Basal medium, supplemented with N2 (1:200), B27 w/o Vitamin A (1:100) (Thermo-scientific) 0.05mM MEM non-essential amino acids (Gibco), L-glutamine (1:100, Gibco), Pen-strep (100 µg/ml each), Insulin (1.6g/L, Sigma Aldrich), 0.05 mM β-Mercaptoethanol (Life Technologies). | 4 |
| 5 | Human brain organoid differentiation medium | 1:1 DMEM/F12 and Neural Basal medium, supplemented with N2 (1:200), B27 w/o Vitamin A (1:100) (Thermo-scientific) 0.05mM MEM non-essential amino acids (Gibco), L-glutamine (1:100, | 20 |
|  |  | Gibco), Pen-strep (100 µg/ml each), Insulin (1.6g/L, Sigma Aldrich), 0.05 mM β-Mercaptoethanol (Life Technologies), and supplemented with 5 µM SB431542 (Selleckchem, USA) and 0.5 µM Dorsomorphin (Sigma-Aldrich, USA) |  |
| 6 | Human brain organoid maturation medium | 1:1 DMEM/F12 and Neural Basal medium, supplemented with N2 (1:200), B27 w/o Vitamin A (1:100) (Thermo-scientific) 0.05mM MEM non-essential amino acids (Gibco), L-glutamine (1:100, Gibco), Pen-strep (100 µg/ml each), Insulin (1.6g/L, Sigma Aldrich), 0.05 mM β-Mercapto ethanol (Life Technologies) | ~ 150-180 |

To assess the overall versatility of the Hi-Q approach, we generated organoids from six independent iPSC lines (Four healthy and two derived from microcephaly patients). Notably, in this approach, the organoids grew in size progressively over time **(Figure 1F)** (except organoids derived from microcephaly patients, which are described later). In this Hi-Q platform, we generated 13093 organoids across 37 batches **(Table 2).** Measuring 300 randomly selected Hi-Q brain organoids across four iPSC lines revealed that organoid size was highly consistent within a batch and across iPSC lines **(Figure 1G).** Furthermore, the organoids showed a consistent and proportional size increase from day 20 to 60 across all iPSC lines. This indicates that organoids do not aberrantly vary in growth with our Hi-Q approach **(Figure 1H).** In all cases, the organoids displayed high integrity as we detected only one or two disintegrated organoids in a batch of 300 **(Figure 1I)**. These results suggest that the Hi-Q approach is robust, versatile, and easy to handle.

#### Time-resolved single-cell RNA-sequencing of Hi-Q brain organoids reveal similar cell diversities and are free from ectopic stress-inducing pathways

To dissect the cell diversity of Hi-Q brain organoids and correlate it to the human brain, we performed single-cell RNA-sequencing (scRNA-seq). We analyzed 16,228 cells isolated from three organoids at each time point of day 60, 90, and 150. The sequenced cells belonged to cell types such as proliferating radial glia (Pro-RG) cells expressing TTYH1, intermediate precursor cells (IPC) expressing MKI67 and NUSAP1, inhibitory neurons (IN) expressing GAD2, and excitatory neurons

(EN) expressing NEUROD2 and NEUROD6 **(Figure 2A and S1A-C).** We then applied *the k*-nearest neighbor network (*k*nn) approach to analyze distances and progression of transcriptional changes among cells in the 2D representation. All three stages of organoids comprise structural, immature, proliferating, and neuronal populations **(Figure 2A-C)**. Notably, cells annotated as proliferating radial glia cells (RG) and IPCs correspond to the cells in the S and G2/M phases **(Figure S1D-E)**.

**Figure 2:**
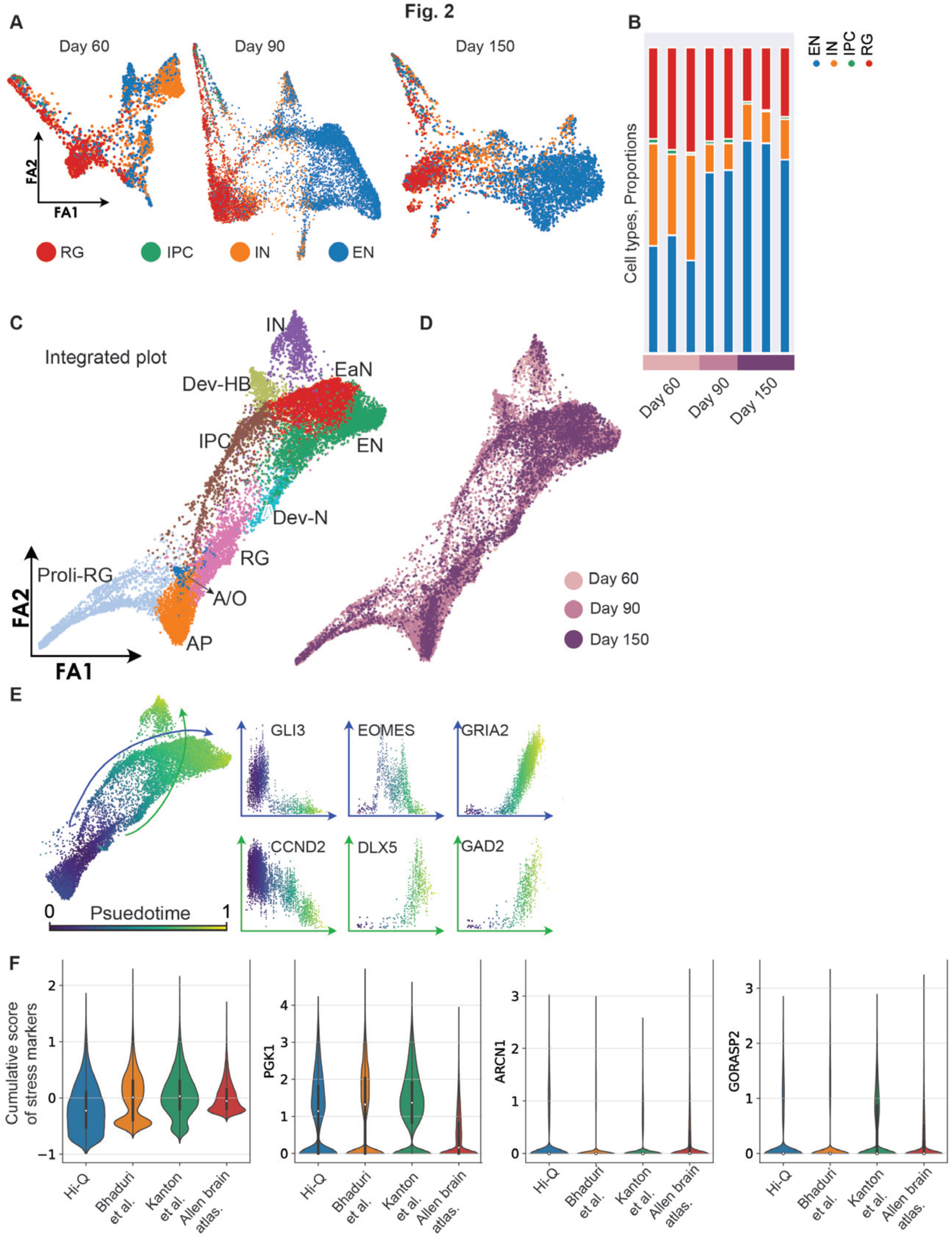
Cell type diversity in Hi-Q brain organoids across maturation. **A.** Force Altas (FA) 2D representation of the neighbor graph of Hi-Q brain organoids at different time points of development (Day 60, 90, and 150). Cells are colored according to the label with the highest score given by the Lasso logistic regression model trained on primary brain data ^40^. **B.** Histogram plot reporting the proportion of different cell types at different time points of development. **C-D.** FA representation of the neighbor graph of the integrated datasets from all stages. In panel **C,** clusters are annotated using the expressed gene markers: Pro-RG, proliferating radial glia; AP, apical progenitors; RG, radial glia; A/O, astrocytes, and oligodendrocytes; IPC, intermediate precursor cells; Dev-N, developing neurons; Dev-HB, developing hindbrain; EaN, early neurons; EN, excitatory neurons; IN, inhibitory neurons. In panel D, cells are colored according to the time they were sampled. **E.** The subset of the whole integrated dataset, except the pro-RG cluster, colored by pseudotime, showing the two trajectories of developing IN (green arrow) and EN (blue arrow). The scatter plots show the expression range of characteristic IN and EN genes for each cell in function of the pseudotime. **F.** Violin plots comparing the level of expression of PGK1, ARCN1, GORASP2, and the cumulative score of the three between Hi-Q brain organoids, other brain organoids ^13^ ^22^ and primary data from the literature.

We then used the annotated cell types to evaluate the degree of similarities and maturation at each stage of Hi-Q brain organoids. First, we noticed that individual organoids showed similar cell proportions at the same age. Next, as the organoids progressively matured, we observed an accumulation of mature neuronal types, especially excitatory neurons (EN), in day 90 and 150 organoids **(Figure 2B)**. We then applied principal component analysis to assess the organoids’ similarities at the transcriptional level within an age group. Importantly, this analysis revealed that the cellular composition of individual organoids exhibited a high similarity with organoids from the same age groups and widely differed from other age groups, indicating that the Hi-Q approach generates highly reproducible organoids and is reliable **(Figure S1F).** Analyzing the cell types computed from the -*k*nn analysis that integrated cell diversities of all three age groups of organoids highlighted the presence of apical neural progenitors (AP), radial glia (RG), proliferating radial glia (Pro-RG), astrocytes and oligodendrocytes (A/O), intermediate precursor cells (IPC), developing neurons (Dev-N), developing hindbrain (Dev-HB), early neurons (EaN), excitatory neurons (EN) and inhibitory neurons (IN) **(Figure 2C-D).**

To further analyze the developmental trajectories of terminally developed populations within the dataset, we used a pseudotime analysis based on diffusion from the RG cluster **(Figure 2E).** The gene expression patterns across the two trajectories agree with the unbiased, calculated markers from the cell types. We observed mainly two trajectories. The first trajectory (blue line) expressing markers of dorsal telencephalon development (GLI3, EOMES, and GRIA2) leads to the formation of ENs. The second trajectory (green line), expressing markers of ventral telencephalon development (CCND2, DLX5, GAD2) ^22^, concludes with the appearance of INs **(Figure 2E and Figure S1G-H).**

A recent work reported that activation of cellular stress pathways in organoids interferes with the developmental process required to generate distinct cell identities of the human brain^13^. Thus, optimizing a method that allows the generation of brain organoids free from cellular stress pathways is critical. To test if Hi-Q brain organoids are free from those ectopic stress-inducing pathways, we analyzed the expression level of previously described stress markers PGK1, ARCN1, and GORASP2 ^13^. Notably, Hi-Q brain organoids showed a lower stress marker expression than published brain organoid datasets but slightly higher than in adult human brain datasets ^13,22^ **(Figure 2F).** In summary, our temporally resolved sc-RNA-sequencing data reveal that the Hi-Q approach can generate brain organoids with reproducible levels of cell diversity free from ectopically induced stress pathways.

#### Hi-Q brain organoids progressively develop over time and reveal mature neuronal markers

Next, we analyzed the cytoarchitecture of the organoids by histology. We selected a panel of markers for each cell type in organoids derived from the IMR90 on day 20 and day 60 and quantified them. Our selected panel included Nestin and SOX2 (progenitors), DCX (early neurons), Acetylated α-tubulin (neurons and cilia), Arl13b (cilia), Actin (neurons), TUJ-1 (pan-neuronal), MAP2 (cortical neurons), CTIP2 (layer 4 and 5-specific), Tau (cortical neurons), PCP4 (Purkinje neurons), GAD67 (glutamatergic neurons), Synapsin-1 (presynaptic), and PSD95 (post-synaptic). To preserve the organoid integrity, we conducted whole-mount immunostaining and confocal imaging of intact organoids after tissue clearing as previously described ^23,24^.

Day 20 organoids mainly exhibited early developmental markers of progenitors and early neurons and barely showed distinct cortical plates decorated by mature neuronal markers. Neuronal markers such as MAP2 and TUJ-1 were expressed but only localized in the cell body and did not entirely segregate into axons, indicating that these organoids are at the early stage of neuronal differentiation **(Figure 3A).** In contrast, day 60 organoids were bigger and exhibited an enriched level of mature neuronal markers with distinguishable cortical plates of similar thickness. Likewise, the proportions of cortical and layer-specific neurons increased with time, with Tau, PCP4, PSD95, Synapsin 1, and CTIP2-positive cells increasing in day 60 organoids **(Figure 3B, Figure S2, and Movie 2).** These data indicate that whole-brain organoids cultured via the Hi-Q approach generate cell types of the early brain and differentiate into mature cell types distinct from the early brain’s germinal zones.

**Figure 3:**
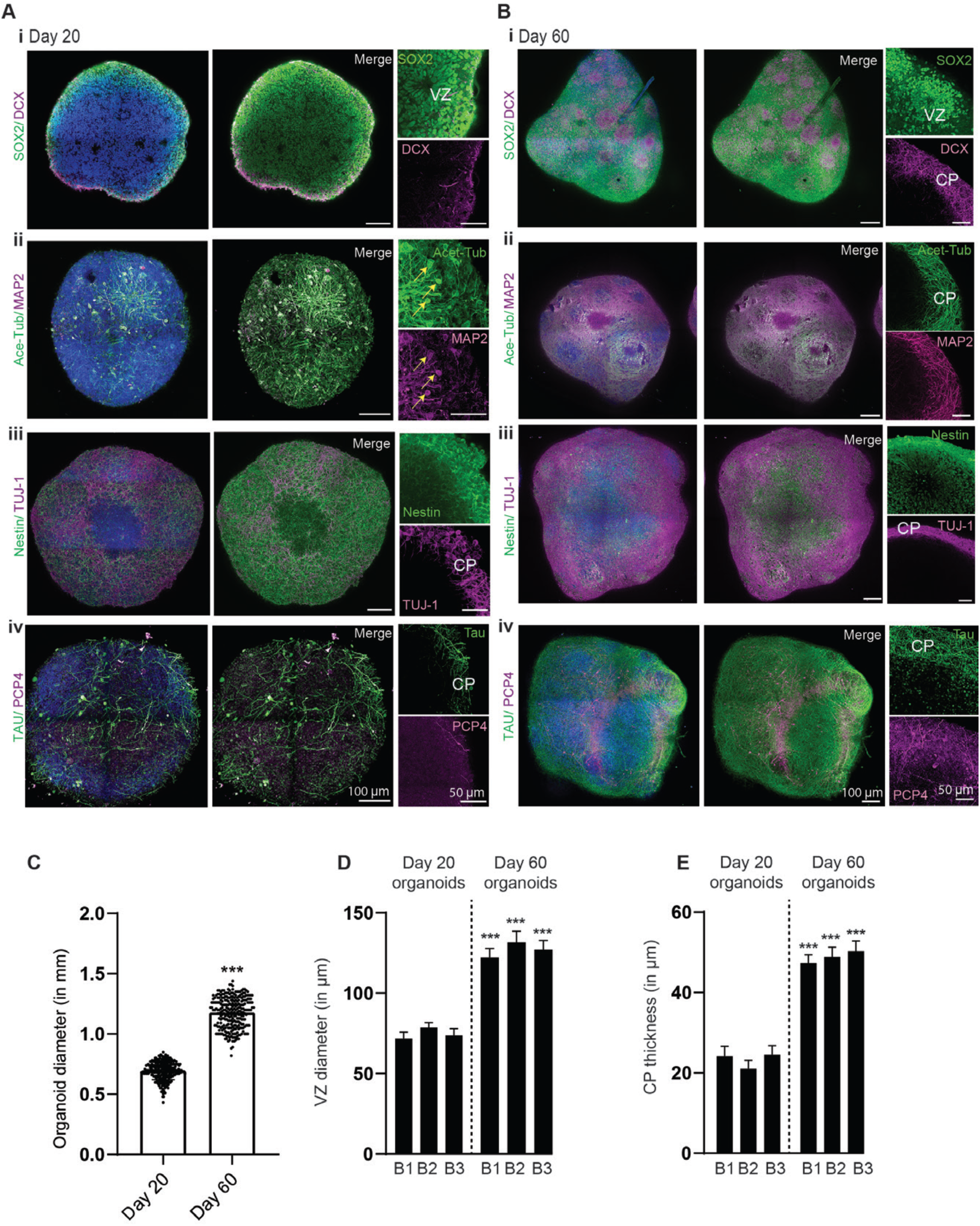
Hi-Q brain organoids mature over time. **A.** Tissue clearing and wholemount staining of day 20 **(Ai-iv)** and 60-day **(Bi-iv)** old organoids show that Hi-Q organoids mature over time. SOX2 and Nestin label NPCs at the ventricular zone (VZ), DCX, Acetylated tubulin, MAP2, TUJ-1, PCP4, and Tau mark neurons at the cortical plate (CP). Note cell body localization of acetylated α-tubulin, MAP2, and TUJ-1 (Arrows in **Ai-iii**) on day 20 organoids are remodeled into defined cortical plates (CP) on day 60 organoids **(Bi-iii).** Likewise, the cortical neuronal markers Tau and PCP4 were remodeled into distinct cortical plates in day 60 Hi-Q brain organoids **(Aiv-Biv)**. Therefore, CPs are primitive or thin on day 20, and organoids thick and distinct on day 60. Representative images are shown, and panels show scale bars. **C-E.** Diagrams quantify size differences between time points **(C)**, VZ diameter **(D),** and CP thickness **(E)**. At least 100 organoids across several batches have been sampled for size comparison. At least 25 independent brain organoids have been sampled for VZ and CP thickness. Statistical analysis was carried out by One-way ANOVA, followed by Tukey’s multiple comparisons test, ***P < 0.001. Data presented as mean ± SEM.

### Spontaneous activity and neuronal networks in Hi-Q brain organoids

Next, we investigated whether Hi-Q brain organoids exhibit functional activity and form active neuronal networks. A reliable indicator for neuronal activity and network formation in the developing and mature brain is intracellular Ca^2+^ signaling ^25^. To probe for functional activity, we performed imaging of intracellular Ca^2+^ in brain organoids at days 30, 40, 50, and 150. To this end, we loaded the organoids with the calcium indicator dye Oregon Green BAPTA1-AM (OGB-1) by bolus injection, staining essentially all cell bodies in the field of view **(Figure 4A)**. Live imaging of OGB-1 fluorescence revealed vivid spontaneous intracellular Ca^2+^ signaling in all preparations and stages analyzed **(Figure 4B)**, with roughly half of all cells (42-54%) being active in the investigated age groups **(Figure 4C)**. Ca^2+^ signals were variable in amplitude and duration. We detected both single transients, lasting 10-20 seconds, and burst-like events, lasting 1-2 minutes, at each stage. Moreover, ultra-slow Ca^2+^ fluctuations were observed, to which faster events usually added on **(Figure 4B)**. The activity was not generally synchronized across the cells in the field of view, although single events occasionally occurred in several cells simultaneously (see asterisks in **Figure 4B**).

**Figure 4:**
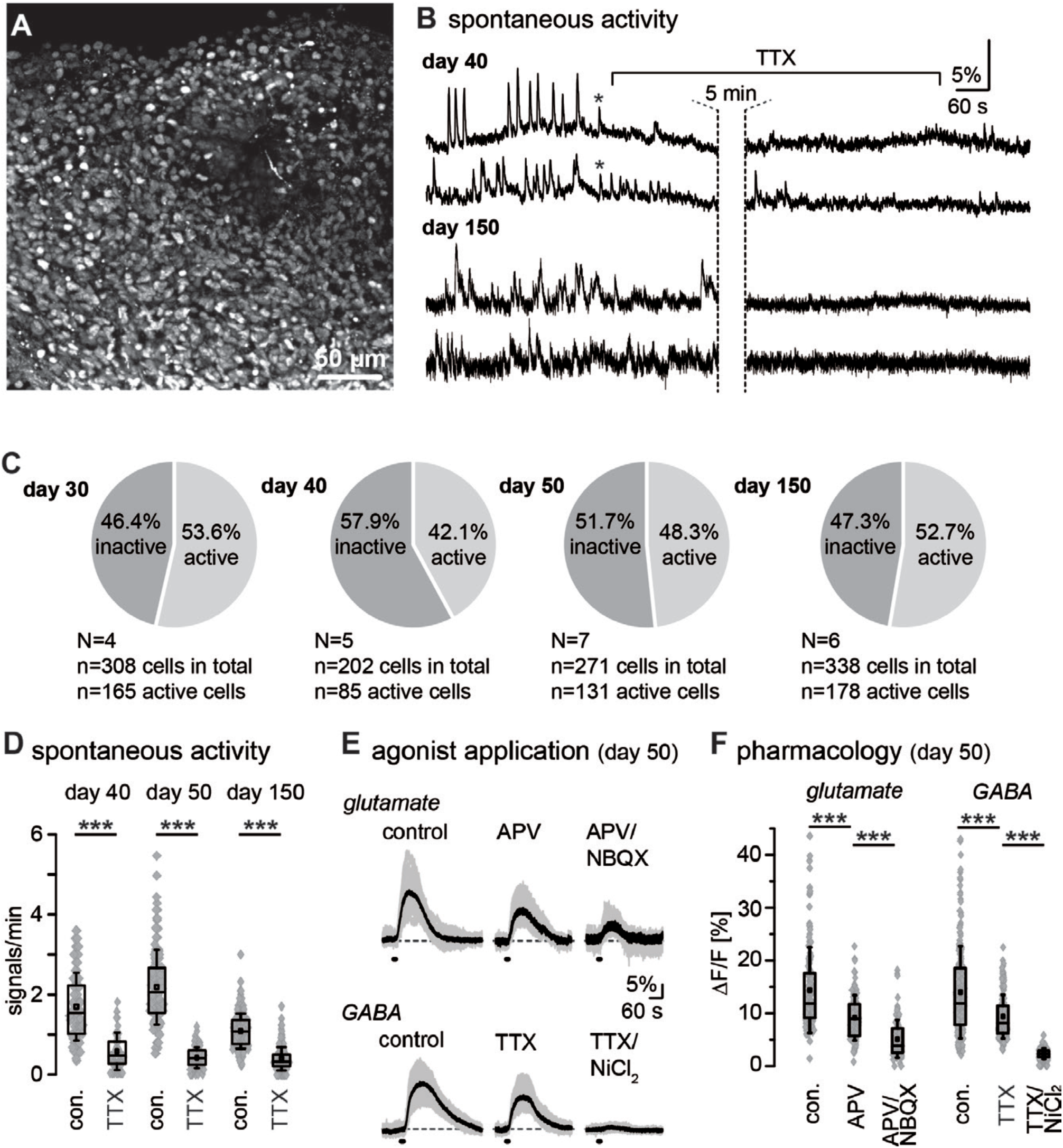
Spontaneous and evoked activity in neural networks in Hi-Q brain organoids. **A.** Fluorescence intensity image of an organoid loaded with the calcium indicator OGB-1. **B.** Exemplary traces of single cells revealing spontaneous calcium signaling in day 40 and day 150 old brain organoids. Asterisks highlight occasional synchronized activity. Wash-in of tetrodotoxin (TTX; 1 µM) strongly dampens spontaneous calcium signaling. **C.** Pie charts illustrating active and inactive cells in brain organoids across all age groups (days 30, 40, 50, and 150). **D.** Box plots showing median (line), mean (square), interquartile range (box), and standard deviation (whiskers) of the frequency of spontaneous calcium activity under control conditions and in the presence of TTX on day 40, 50, and 150 old brain organoids. Grey diamonds represent single data points/cells analyzed. **E.** Intracellular calcium transients evoked by bath application of 1 mM glutamate (upper row) or 1 mM GABA (lower row) under control conditions and in the presence of ionotropic glutamate receptor inhibitors (APV for NMDARs and NBQX for AMPARs) or the presence of inhibitors of voltage-ion channels (TTX for Nav and NiCl2 for Cav) in day 50 brain organoids. Grey traces show individual cellular responses in one particular experiment; black traces are averages of the individual traces. **F.** Box plots showing median (line), mean (square), interquartile range (box), and standard deviation (whiskers) glutamate- or GABA-induced calcium responses of day 50 brain organoids in control conditions and in the presence of indicated receptor/channel blockers. Grey diamonds are single data points/cells analyzed.

In organoids on days 40, 50, and 150, spontaneous activity was strongly dampened (p=1.39E-14; p=7.96E-34; p=3.88E-48) in the presence of tetrodotoxin (TTX; 1 µM) that blocks voltage-gated Na^+^ channels, suggesting that Ca^2+^ signals were mostly secondary to action potential generation and activation of voltage-dependent Ca^2+^ channels **(Figure 4D)**. Next, we applied the neurotransmitters glutamate and GABA by bath perfusion for 10 seconds to further characterize functional activity in brain organoids. Notably, in day 40, day 50, and 150-day-old organoids, all cells tested responded to the application of 1 mM glutamate (day 40: N=3, n=200; day 50: N=3, n=142; day 150: N=3, n=230) or 1 mM GABA (day 40: N=3, n=173; day 50: N=3, n=211; day 150: N=3, n=218) with large and long-lasting Ca^2+^ transients **(Figure 4E, F)**. Glutamate-induced Ca^2+^ signals at day 50 were significantly reduced by the application of blockers of the NMDA-receptor blocker APV (100 µM) (N=3, n=133; p=2.84E-12), and further dampened by combined application of APV with the AMPA-receptor blocker NBQX (50 µM) (N=3, n=108; p=3.50E-29) **(Figure 4E-F).** Ca^2+^ transients induced by GABA application were significantly reduced upon inhibition of action potential generation by TTX (N=3, n=204; p=3.85E-14), and nearly completely suppressed by additional perfusion with NiCl2. Taken together, these results indicate that Hi-Q brain organoids develop functionally active neural networks with cells expressing voltage-gated, TTX-sensitive Na^+^ channels and voltage-gated Ca^2+^ channels, major hallmarks of differentiated neurons. Moreover, cells respond to glutamate and GABA, the brain’s primary neurotransmitters, respectively. Using pharmacological tools, we also obtained evidence for functional expression of ionotropic glutamate receptors, namely AMPA- and NMDA receptors. Finally, the sensitivity of GABA-induced Ca^2+^ signals to TTX and NiCl2 indicates that Ca^2+^ transients evoked by bath application GABA were secondary to cellular depolarization upon activation of ionotropic GABAA receptors ^26^ ^27^.

#### Hi-Q brain organoids can be cryopreserved, thawed, and re-cultured

Unlike patient-derived liver and intestine organoids, brain organoids have not been cryopreserved, thawed, and re-cultured, an aspect that limits the flexible and economical use of brain organoids ^28, 29^. We could successfully freeze 8-day-old Hi-Q brain organoids, cryopreserved them in liquid nitrogen for four days and re-cultured them after thawing **(Figure 5A, see method section).** Notably, 8-day-old organoids already exhibit TUJ-1-positive primitive cortical plate and SOX-2-positive proliferating neural progenitor cells (NPCs). To re-culture the organoids, we first thawed them into the organoid culturing medium for 48h in a stationary petri dish and then cultured them in spinner flasks for several days **(Figure 5B).** Although 48 hours post-thawing, organoids did appear slightly deformed, twelve days after re-culturing (20-day-old organoids), the organoids were morphologically similar to control organoids that had never been frozen. Notably, thawing and re-culturing did not cause a significant disassociation or change in size **(Figure 5C-D).** Importantly, we could also thaw and culture organoids in a petri dish without involving a spinner flask. Although the initial growth rate of thawed organoids was slower than controls, they followed a similar growth profile to control organoids **(Figure S3A).** In contrast to 8-day-old organoids, we could not successfully re-culture day 20 and day 30-old brain organoids frozen similarly (data not shown).

**Figure 5:**
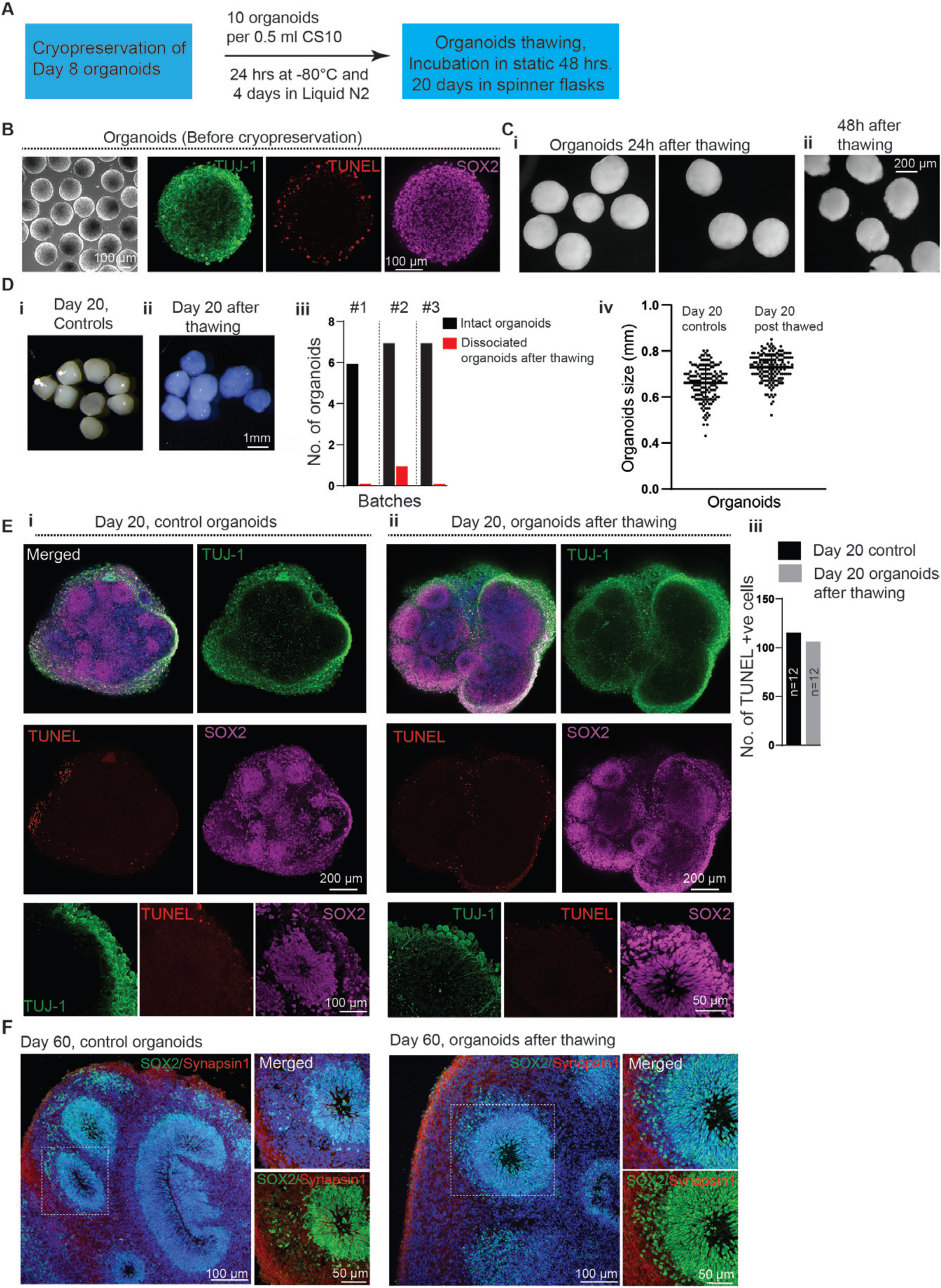
Cryopreservation, thawing, and re-culturing of Hi-Q brain organoids. **A.** A schematic summarizes the cryopreservation and freeze-thawing of Hi-Q brain organoids. **B.** Eight-day-old organoids before cryopreservation. TUJ-1 marks early neurons (green), TUNEL (red) shows the dead cells at the periphery, and SOX2 labels NPCs (magenta). Panels show the scale bar. **C.** Hi-Q organoids 24 hours after thawing **(i)** are still intact. After 48 hours of thawing **(ii),** organoids show a slightly deformed edge. The panel shows the scale bar. **D.** Day 20 Hi-Q organoids that have never been cryopreserved **(i).** The right panel **(ii)** shows a group of organoids 20 days after thawing. The color differences are due to different microscopes with filter settings. The panel shows the scale bar. The Bar graph **(iii)** shows fractions of intact (black) and disintegrated (red) organoids after thawing across three independent batches. **(iv)** At least eighty (n=80) organoids from three independent batches (N=3) were thawed and analyzed. One-way ANOVA carried out statistical analysis. There are no significant differences in the size of Hi-Q brain organoids between controls (never frozen) and freeze-thawed. Student‘s t-test carried out statistical analysis. **E-F.** Comparison of the cytoarchitecture of day-20 (E) Hi-Q brain organoid (control, never frozen) **(i)** and frozen and thawed organoid **(ii).** Magnified panels at the bottom show a ventricular zone (VZ) with its typical cytoarchitecture of early neurons (primitive cortical plate, green marked by TUJ-1) and proliferating NPCs (magenta marked by SOX2). TUNEL (red) labels dead cells; there is no difference between control and freeze-thawed organoids **(iii).** Panel **F** shows the cytoarchitecture of 60-day-old organoids stained with SOX2 (green) and Synapsin1 (red). Panels show scale bars. At least two (n=12) randomly chosen organoids across two independent batches (N=2) were compared to Day 20 controls. Student ‘s t-test carried out statistical analysis.

We then analyzed the integrity and composition of cytoarchitectures after thawing and re-culturing. We immunostained them for SOX2, TUJ-1, and TUNEL, which labels NPCs, primitive cortical plate, and dead cells, respectively. Our analysis revealed that the re-cultured organoids exhibit strikingly similar cytoarchitectures to age-matched control groups, which were not cryopreserved and re-cultured. Notably, there was no difference in the frequencies of TUNEL-positive cells **(Figure 5E, S3, and Movie 3).** By day 60, thawed and re-cultured organoids displayed synapsin-1-positive mature neuronal types and architectures, which were indistinguishable to control organoids, which were never cryopreserved **(Figure 5F).** These data indicate that organoids generated with the Hi-Q platform are amenable to cryopreservation and are viable and healthy after thawing and re-culturing.

#### Patient-derived Hi-Q brain organoids can recapitulate distinct forms of neurodevelopmental disorders

We aimed to model two distinct neurodevelopmental disorders to test Hi-Q brain organoids’ versatility in disease modeling. Firstly, we analyzed a primary microcephaly condition due to a loss-of-function homozygous mutation in a centrosome protein CDK5RAP2. To generate primary microcephaly iPSCs, we reprogrammed patient skin fibroblasts carrying a homozygous mutation in CDK5RAP2 ^30^ **(Figure S4A-C).** Second, we modeled progeria-associated Cockayne syndrome using iPSCs derived from a patient exhibiting Cockayne syndrome (also known as Cockayne syndrome B, CSB). CSB exhibits severe neurological defects caused by a mutation in ERCC6 (also known as Cockayne syndrome B, CSB). CSB is a protein implicated in the transcription-coupled nucleotide excision repair pathway in DNA damage ^31–33^.

Although we used the same number of iPSCs (Healthy control, CDK5RAP2-mutated, and Cockayne syndrome) to generate Hi-Q brain organoids, age-matched CDK5RAP2 patient-derived organoids (Hereafter CDK5RAP2 organoids) were smaller in size. On the other hand, organoids generated from CSB patient-derived iPSCs (hereafter CSB-Organoids) were more prominent in size, indicating that cellular functions leading to neural epithelia formation are abnormal in both models **(Figure 6A)**. In addition, immunostaining for SOX2-positive progenitors and TUJ1-positive early neurons on 20-day-old organoids revealed that control organoids displayed structurally organized ventricular zones (VZ) abundant with progenitors in a compact organization and a layer of neurons forming a primitive cortical plate **(Figure 6Bi).** In contrast, CDK5RAP2 organoids revealed structurally compromised VZs with reduced diameter, less compact progenitors’ organization, and a dispersed cortical plate with TUJ1-positive neurons spreading through organoids **(Figure 6Bii).**

**Figure 6:**
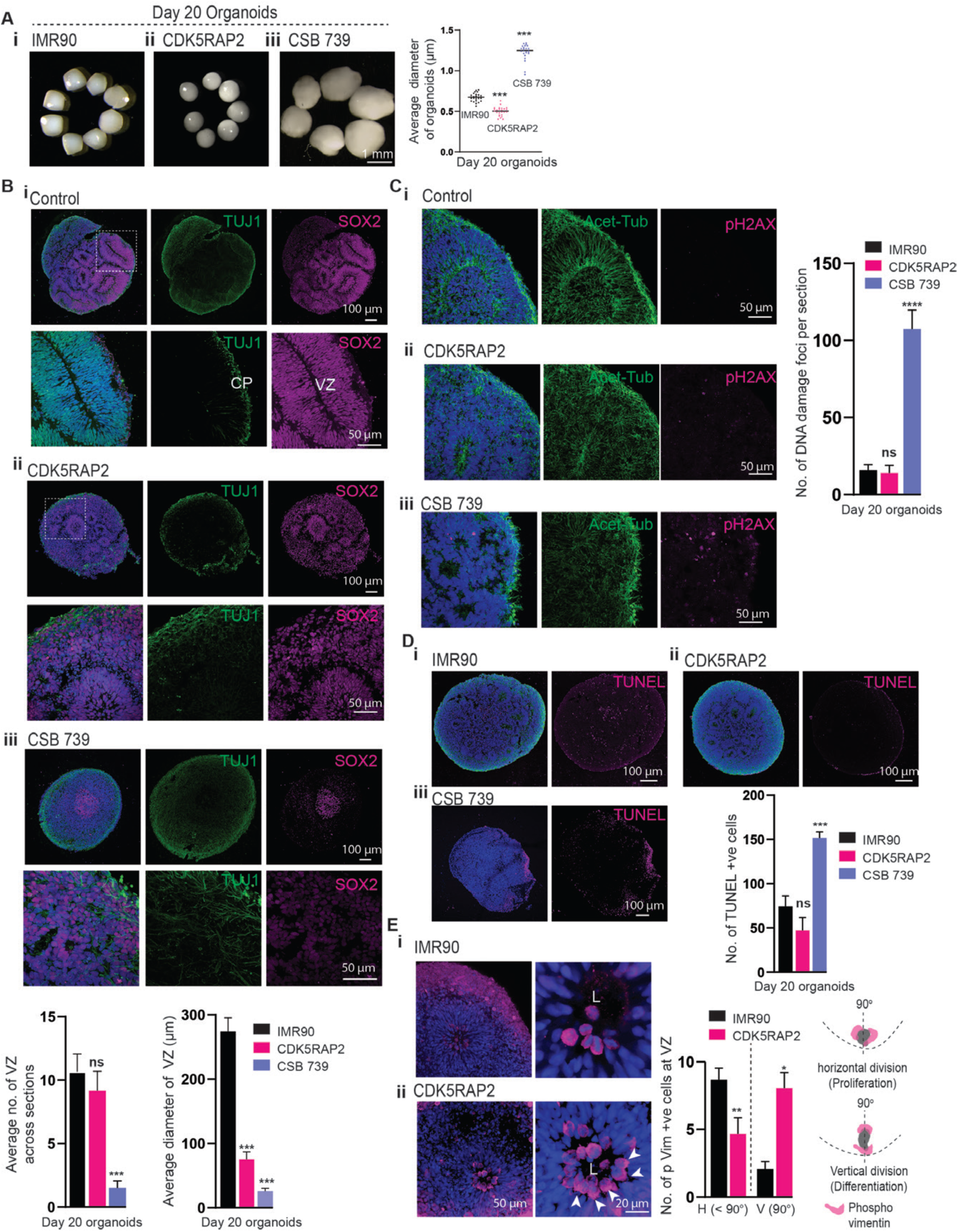
Hi-Q brain organoids model microcephaly and brain organization defects. **A.** Comparison of healthy control (IMR90) and Hi-Q brain organoids derived from mutant patients harboring mutations in CDK5RAP2 and Cockayne syndrome B gene (CSB 739) **(i-iii).** CDK5RAP2 brain organoids **(ii)** are microcephalic as they are significantly smaller than healthy organoids. On the other hand, CSB organoids are significantly larger than healthy controls. The panel shows the scale bar. At least twenty (n=20) randomly chosen day 20 organoids were analyzed across three independent batches (N=3). Statistical analysis was carried out by One-way ANOVA, followed by Tukey’s multiple comparisons test, ***P < 0.001. Data presented as mean ± SEM. **B.** Tissue sections of healthy control (IMR90), CDK5RAP2, and CSB 739 brain organoids **(i-iii)**. The magnified panel under each variety shows VZ. SOX2 labels NPCs, and TUJ-1 marks primitive cortical plate (CP). Compared to healthy organoids (IMR90), CDK5RAP2 organoids have slightly disorganized and smaller VZ. Bar graphs at the bottom quantify the average number of VZs and their diameter in each kind. Panels show the scale bar. At least twenty sections from three different organoids of each group were analyzed. Statistical analysis was carried out by One-way ANOVA, followed by Tukey’s multiple comparisons test, ***P < 0.001. Data presented as mean ± SEM. **C.** CSB 739 **(iii)** but not healthy **(i)** or CDK5RAP2 **(ii)** organoids display a significantly increased level of pH2AX-positive nuclei (magenta) indicative of DNA double-stranded breaks. The bar graph at the right quantifies the relative proportions of pH2AX-positive nuclei. Panels show the scale bar. An average of at least twenty sections from three different organoids of each group was considered. Statistical analysis was carried out by One-way ANOVA, followed by Tukey’s multiple comparisons test, ***P < 0.001. Data presented as mean ± SEM. **D.** CSB 739 **(iii)** but not healthy **(i)** or CDK5RAP2 **(ii)** organoids display a significantly increased level of TUNNEL-positive cells (magenta) indicative of dead cells. The bar graph quantifies the relative proportions of TUNNEL-positive nuclei. An average of at least twenty sections from three different organoids of each group was considered. Statistical analysis was carried out by One-way ANOVA, followed by Tukey’s multiple comparisons test, ***P < 0.001. Data presented as mean ± SEM. Panels show the scale bar. **E.** Hi-Q brain organoids reveal the kinetics of the apical progenitor division plane. P-Vimentin selectively labels the dividing apical progenitors at the VZ’s apical side. Healthy brain organoids (IMR90) predominantly display dividing progenitors whose division plane is horizontal to the VZ lumen **(i)**. In contrast, CDK5RAP2 brain organoids **(ii)** harbor apical progenitors whose division plane is mainly vertical to the VZ lumen. The bar diagram quantifies the distribution of the division plane. H, horizontal. V, vertical. A schematic at the right shows horizontal and vertical division planes. An average of at least fifteen sections from three different organoids of each group was considered. Statistical analysis was carried out by One-way ANOVA, followed by Tukey’s multiple comparisons test, *P < 0.1. Data presented as mean ± SEM. Panels show the scale bar.

CSB organoids (CSB 739), on the other hand, did not exhibit recognizable VZs, and the progenitors were less densely packed and randomly distributed with weakly positive SOX2. Moreover, TUJ-1 positive neurons did not form a distinct cortical plate; instead, they were broadly diffused in the organoid tissue, suggesting that brain organization is perturbed in CSB organoids **(Figure 6Biii and S4D-F)**. Finally, turning our analysis for spontaneous DNA damage and apoptotic cell deaths, we noticed that CSB-organoids harbored many pH2AX-positive and TUNEL-positive cells **(Figure 6C-D).** These findings revealed that without functional CSB, neural tissues undergo extensive DNA damage, thereby perturbing brain organization.

Notably, neither DNA damage nor cell death appears to cause microcephaly in CDK5RAP2 organoids, as we did not notice a significant increase in pH2AX-positive cells and TUNEL-positive cells **(Figure 6C-D)**. Thus, we reasoned that the microcephalic phenotype of brain organoids could be due to defective NPCs’ proliferation, leading to premature differentiation of NPCs and NPCs loss. To test this, we analyzed the kinetics of NPCs proliferation by analyzing the division plane of p-Vimentin-positive apical progenitors that form the lumen of the VZ. In contrast to controls, which harbored increased frequencies of horizontally oriented mitotic NPCs, we identified that patient-derived organoids harbored mostly vertically oriented mitotic NPCs. This indicated an unscheduled switching of the division plane, leading to premature differentiation with depletion of progenitors **(Figure 6E).** These results explain that the loss of CDK5RAP2 perturbs the horizontal orientation of the spindle in the patient tissues that are crucial for the symmetric expansion of the NPCs pool. These comparative analyses, in summary, indicate that the Hi-Q brain organoids are robust in modeling neurodevelopmental disorders.

#### Hi-Q brain organoids model glioma invasion and are amenable to medium-throughput compound screening

Brain organoids hold promise for 3D-organoid-based compound screening. However, this requires generating a high quantity of organoids with increased homogeneity. As Hi-Q brain organoids satisfy this prerequisite, we attempted to develop a proof-of-principle experiment adapting Hi-Q brain organoids for a medium-throughput drug screen. Brain organoids have recently been used to reveal the neuro-invasion behavior of glioblastoma (GBM) ^24,34^. GBM harbor glioma stem cells (GSCs) that infiltrate the brain and account for the fatal nature of this disease for which there is no promising cure^35^. Thus, we investigated whether brain organoids can be used to identify compounds inhibiting the GSC invasion.

First, we set out to determine the invasion behavior of glioblastoma (GBM) in Hi-Q brain organoids. We labeled a patient-derived GSC line (#450) with mCherry and adapted our recently established brain organoid-based GBM invasion assay to Hi-Q brain organoids ^23,24^. Importantly, Hi-Q brain organoids could faithfully recapitulate the invasive behavior of GSCs. In brief, when 5000 suspended GSCs or GSC spheroids were applied to day-40 Hi-Q brain organoids, GSCs adhered to the surrounding organoids within 24 hours. Within 24-72 hours, GSCs infiltrated the organoids, exhibiting typical in vivo glioma invasion characteristics, such as protrusions with extended microtubes ^36^**(Figure 7A-B)**. Using this setup, we screened compounds that could prevent the neuro-invasion property of GSCs.

**Figure 7:**
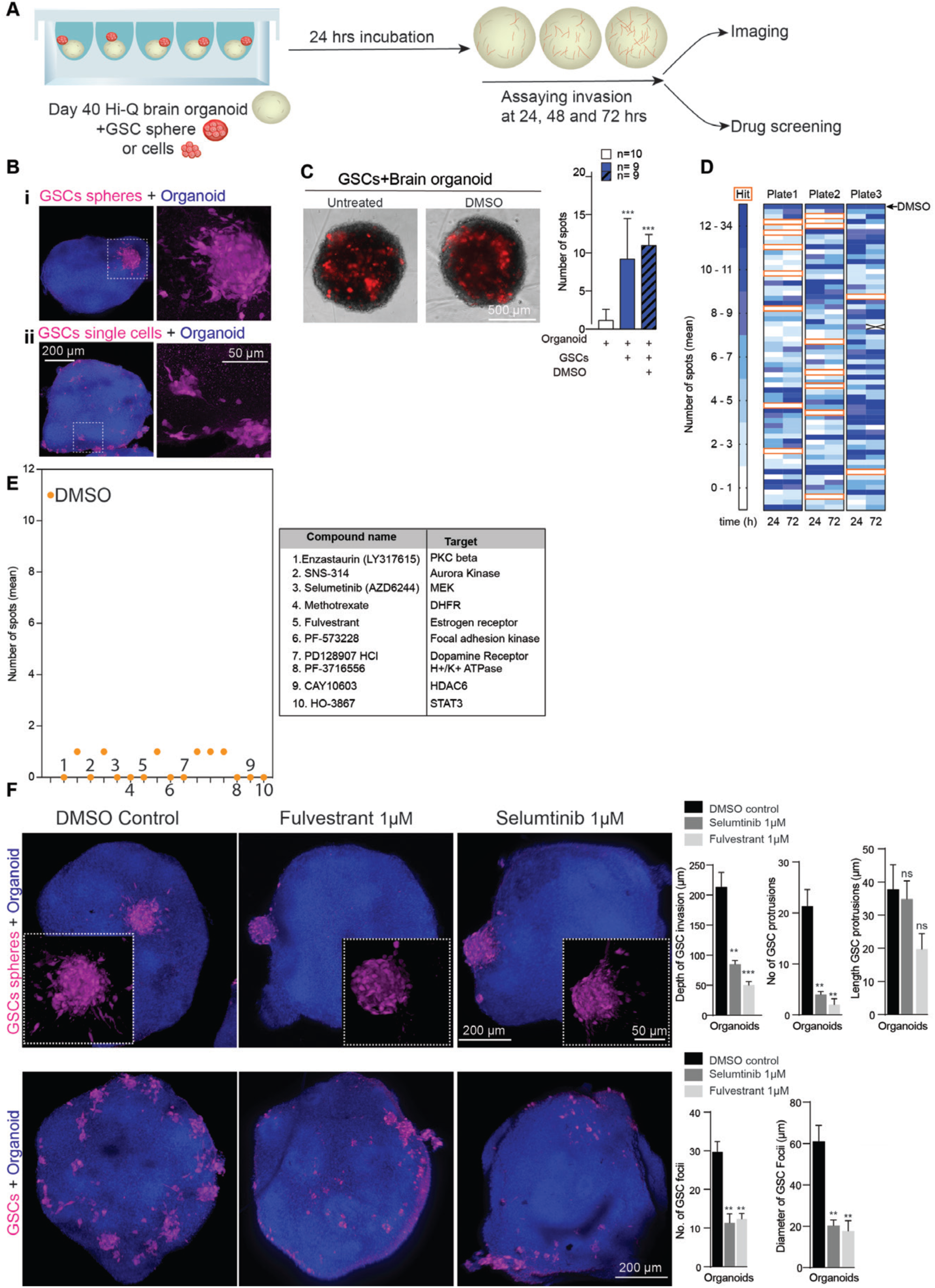
Hi-Q brain organoids allow glioma invasion assays and can adapt medium-throughput drug screening assays to identify anti-glioma compounds. **A.** Experimental scheme for a glioma invasion assay using Hi-Q brain organoids that can be adapted for drug screening. **B.** Hi-Q brain organoids allow GSC invasion as spheres **(i)** or single cells **(ii).** The magnified panel at the right shows invading GSCs with their typical protrusions into the organoid tissues (arrows). The bar diagram at the right quantifies the depth invaded by GSCs. Panels show the scale bar. **C-D.** Assay design adapted to screen the compounds preventing GSC invasion. Representative images showing invading GSC into Hi-Q brain organoids **(C).** At least nine organoids from three independent experiments were analyzed. Statistical analysis was carried out by One-way ANOVA, followed by Tukey’s multiple comparisons test, ***P < 0.001. Data presented as mean ± SEM. Automated imaging and computer-based counting of GSC spots in each organoid. Two organoids were tested for each compound. The algorithm assigns a space when there are no invading GSCs. Dark blue denotes more GSC spots in organoids. Each box represents a well supplied with a compound **(D)**. **E.** Dot plot shows the selected compounds that prevent GSC invasion into organoids. Each dot denotes the number of GSC spots in organoids. The table at the right lists selected ten compounds that can stop the GSCs’ invasions into brain organoids. The biological functions of each compound are also listed. **F.** Quantitative 3D imaging of GSCs invasion into brain organoids. The top panel shows the GSC invasion from spheres. The bottom panel shows GSCs invasions from GSCs suspension. Fulvestrant and Selumtinib significantly prevent GSC invasion into brain organoids as selected compounds. The invaded GSC (as spheres or single cells) were marked with red arrowheads. White arrowheads point to the spheres or cells that fail to invade the organoids in the presence of drugs. Panels show the scale bar. The bar diagram quantifies the inhibitory effects of the selected compounds on GSC invasion. Average measurements from at least six organoids per drug condition were considered. Statistical analysis was carried out by One-way ANOVA, followed by Tukey’s multiple comparisons test, ***P < 0.001, **P < 0.01. Data presented as mean ± SEM).

We consolidated a library of 180 compounds with known biological targets to design a medium-throughput assay **(Table 3).** We began the preliminary screening assay in 96-well format by incubating Hi-Q brain organoids and GSCs for 72 hours with 5 µM of each compound. We imaged each invasion sample over three days with two image acquisition time points in an automated microscope programmed to acquire images at twenty-four and seventy-two hours. We used high-content image analysis to determine the invasion of GSCs into organoids. Here, spots were detected within the organoid region, and their numbers were quantified (see methods section) **(Figure 7C).** Via this assay, we identified sixteen compounds that negatively affected GSC invasion **(Figure 7D-E).** As a secondary screen, we tested the ability of ten of these sixteen compounds (which exhibited the most inhibitory effect) to perturb GSC invasion at 1µM concentration. This assay relied on uniform-sized organoids for comparison. This analysis identified Selumetinib and Fulvestrant as effective inhibitors of GSC invasion into brain organoids as judged by the differences in the number of GSC foci between twenty-four and seventy-two hours of invasion assay **(Figure S5, parts 1 and 2, red dotted box in part 2)**. Selumetinib is a mitogen-activated protein kinase 1, and 2 inhibitors used to treat neurofibromatosis ^37^. Fulvestrant is a selective estrogen receptor degrader (SERD) used to treat advanced breast cancer ^38^.

We then tested whether Selumetinib and Fulvestrant compounds could perturb GSC invasion when supplied as single cells or spheres in Hi-Q brain organoids **(Figure S6 and S7, Low resolution).** We then applied high-resolution quantitative 3D imaging and evaluated the abilities of these compounds in perturbing GSC invasion. Compared to vehicle control, these compounds significantly prevented the invading power of GSCs when supplied with either compact spheres or single cells **(Figure 7F).**

To test whether the drugs identified in our H-Q brain organoid had a significant impact *in vivo*, we grafted GFP-tagged GSC lines onto the striatum of NOD-SCID mice (line GSC#1 and GSC#472. **Figures S8** and **S9**, respectively). After a week of grafting, we treated the animals for three weeks with saline or a combined 20 mg/Kg of Selumetinib and Fulvestrant. In untreated mice, both GSC lines grow extensively at the injected striatum and spread to the white matter paths, like the corpus callosum, optic tract, anterior commissure, and cerebrospinal fluid (CSF) pathways. We imaged histological sections through the tumor epicenter to assess the invasion of the white matter and CSF paths. We analyzed the number of GSCs invading the corpus callosum, optic tract, and the walls of the ventricles in saline and drug-treated subjects **(Figures S8 and S9)**. The drug treatment significantly reduced the tumor volumes of GSCs in the striatum **(Figure S8A and S9A)**. Besides, the drug treatment also significantly reduced the spread of tumor spheres onto the ventricular walls and impaired the invasion of GSCs into the corpus callosum and optic tract **(Figure S8D-E)**. Finally, we analyzed the GSC proliferation using Ki67 immunostaining on sections. We noticed that the drug treatment has significantly reduced the GSC proliferation as we could observe reduced frequencies of cells positive for Ki67 **(Figure S8E).**

In summary, the brain organoids generated *via* the Hi-Q approach are amenable to model glioma and can ultimately serve as a test system to screen potential therapeutic compounds to inhibit the invasive behavior of GSCs.

## Discussion

Brain organoids have shown promise in modeling human brain development and neurological diseases. However, the organoids’ utility has been limited by several shortcomings related to their homogeneity and reproducibility and the difficulty of using them in drug screening settings. Considerable efforts have been invested in developing culturing conditions to ensure high reproducibility across individual brain organoids within a batch ^7^. For example, cortical spheroids have been generated from iPSCs free from the extracellular matrix ^39^. Yet, challenges remain. Notably, a comprehensive study revealed that brain organoids generated from current methods have altered neuronal diversity due to induced stress pathways ^13^, and thus, cannot reliably model diseases. To date, no standardized method is available to fulfill these pitfalls. Yet, developing a brain organoid culturing method that can robustly model various diseases and adapt drug screening strategies is critically needed.

All of these aspects have prompted us to optimize culturing conditions, which led us to generate Hi-Q brain organoids. Notably, our protocol generates large quantities of organoids per batch exhibiting a high degree of similarities in size, shape, cell diversities, and, importantly, free from excessive ectopic stress-inducing pathways **(Figure 2F)**. The Hi-Q approach is robust as we could generate organoids from at least six genetic backgrounds **(Figures 1 and 6).** In addition, the presence of active neuronal networks and increasing cell diversities over time indicate that Hi-Q brain organoids are physiologically relevant **(Figure 4).** Furthermore, we were successful at cryopreserving brain organoids generated with the Hi-Q approach, offering an economic advantage when adapting drug testing and applying biochemical experiments **(Figure 5).**

The Hi-Q approach significantly differs from the current methodology in at least two aspects. Firstly, we avoid embryoid body formation and differentiate iPSC directly into neurospheres using a micropatterned plate. This allowed us to perform the initial differentiation steps of iPSCs in Matrigel and ROCK inhibitor-free conditions and obtain uniform-sized neurospheres, which are the early intermediates of neural tissues. Secondly, the entire protocol follows unguided differentiation and requires minimal manual handling of cells. As the method already regulates the uniform-sized neurospheres, aberrant growth of organoids at later stages with varying sizes is limited.

3D brain organoids recapitulate human brain development principles, and thereby hold promise to decipher neurodevelopmental abnormalities. Thus, we tested the ability of Hi-Q brain organoids to model at least two different types of microcephaly. When modeling CDK5RAP2-mutated microcephaly, Hi-Q brain organoids could recapitulate the premature differentiation of NPCs, a mechanism attributed to causing the depletion of NPCs causing microcephaly **(Figure 6).** Conversely, when modeling progeria-related microcephaly due to mutations in CSB, the Hi-Q brain organoids could display distinct phenotypes exhibiting DNA damage and defects in brain organization. These findings suggest that Hi-Q brain organoids can be adapted to model a variety of neurodevelopmental disorders. Besides modeling neurodevelopmental disorders, we also show that Hi-Q brain organoids apply to the study of glioma invasion behaviors **(Figure 7).**

Although brain organoids hold promise for conducting drug screening studies, one of the most challenging aspects of brain organoid research is to generate a large number of organoids with minimum or no inter-organoid differences within and across batches. Our proof-of-principle experiment successfully used Hi-Q brain organoids in a medium-sized drug screening assay. Using machine learning algorithms, we identified Selumetinib and Fulvestrant compounds that effectively perturbed the invasion behavior of patient-derived glioma stem cells (GSCs) in vitro and mouse xenografts **(Figure 7 and Figure S8-S9).**

In summary, our Hi-Q technology solves many of the limitations in the field of brain organoid research. By generating brain organoids in large quantities in a versatile and robust way, we trust that our approach will pave the path for personalized medicine through disease modeling and high throughput compound screening.

## Acknowledgment

We thank the members of the Laboratory of Centrosome and Cytoskeleton Biology (CCB). JG acknowledges support from the Deutsche Krebshilfe (70114276), Innovation funding from BMBF VIP+ (03VP10540) and Deutsche Forschungsgemeinschaft (DFG, German Research Foundation) – Project-ID 503306912 – FOR5547; R.P. is supported by AIRC (IG 2019 Id.23154); VB is supported by Volkswagen Foundation (Freigeist—A110720) and the Deutsche Forschungsgemeinschaft (BU 2974/3-2 - SPP2127, EXC-2151-390873048-Cluster of Excellence— ImmunoSensation2 at the University of Bonn). CRR: supported by the Deutsche Forschungsgemeinschaft, RU 2795 (“Synapses under Stress”), Ro2327/13-2.

## Resource availability

### Lead contact

Further information and requests for resources and reagents should be directed to and fulfilled by the lead contact, Jay Gopalakrishnan.

### Materials availability

All reagents generated in this study are available from the lead contact with a completed Materials Transfer Agreement.

### Data and code availability

All original code for scRNA analysis of Hi-Q brain organoids has been deposited and is publicly available under the following link: https://github.com/Gpasquini/HiQ_analysis_reproducibility

The lead contact will share all data reported in this paper upon request. Any additional information required to reanalyze the data reported in this paper is available from the lead contact upon request.

### iPSCs details

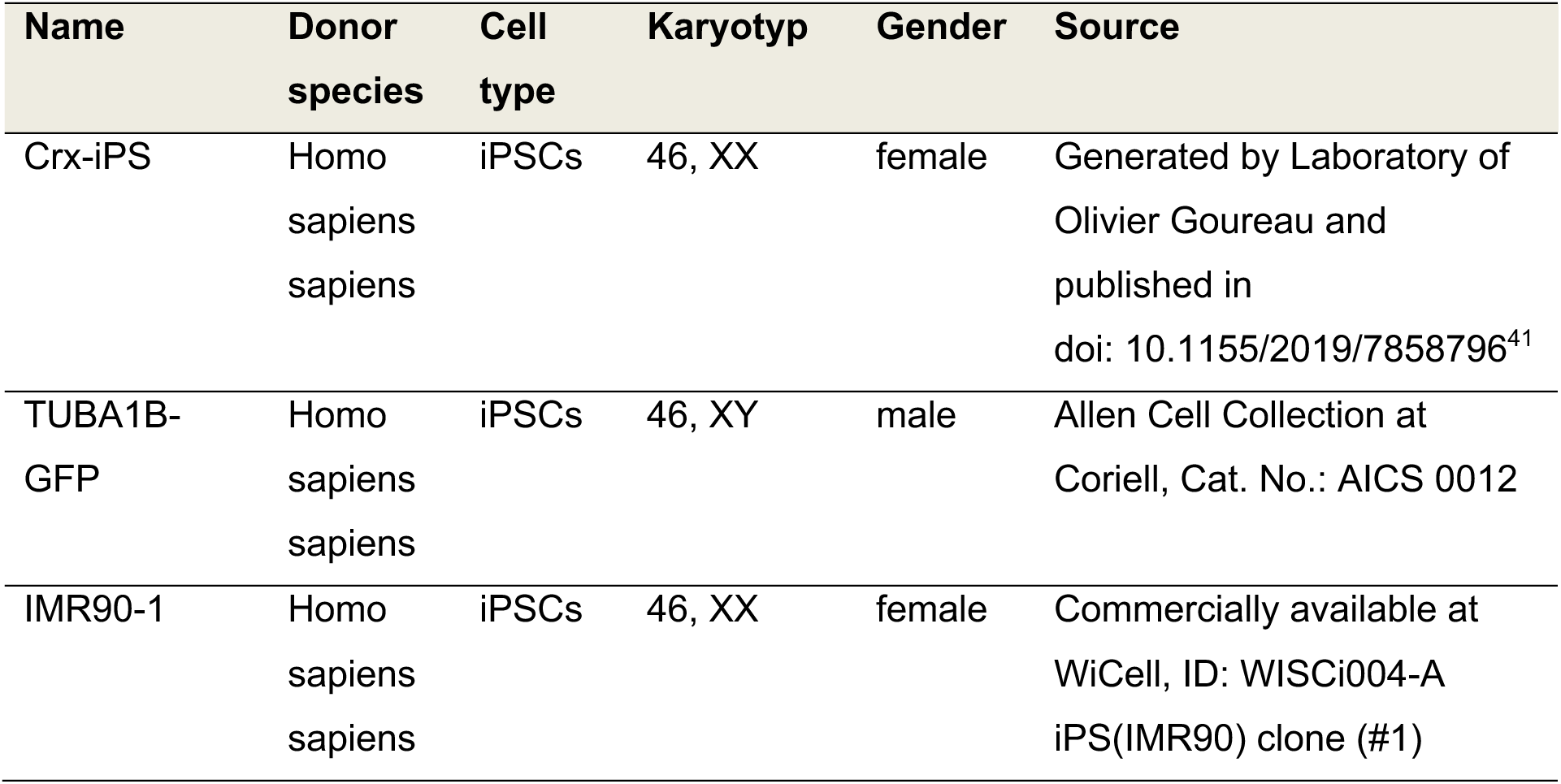
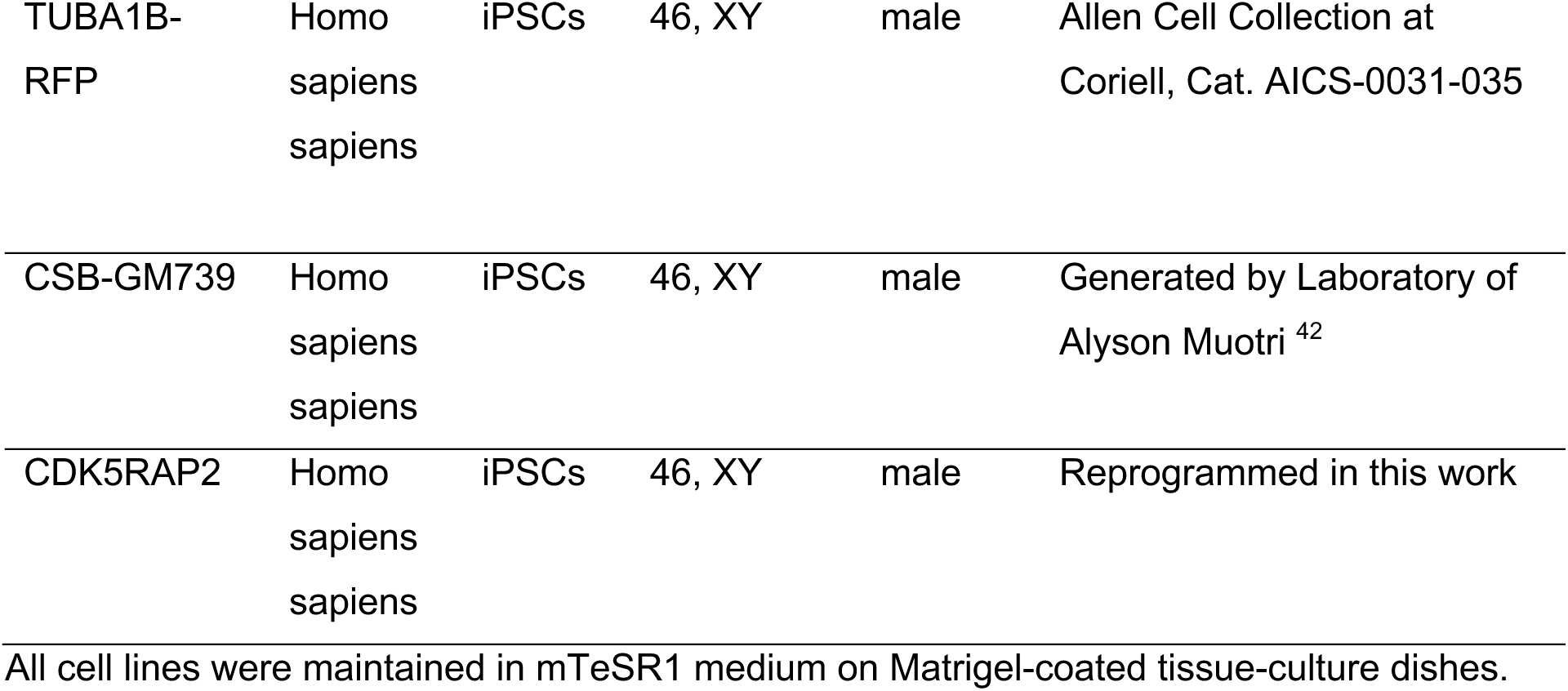

Human iPSCs were cultured in mTeSR1 medium (Stem cell technologies) on Matrigel (Corning) coated cell culture dishes at 37°C and 5% CO2 and routinely checked for mycoplasma contamination with the MycoAlert Kit (Lonza). Media change was carried out once in 3 days. The cells were split 1:2 onto fresh Matrigel-coated dishes using an enzyme-free method, employing ReLSR (Stemcell Technologies, Catalog # 05872) as per the manufacturer’s protocol.

### Generation of Hi-Q brain organoids

#### Seeding iPSCs and neural induction

iPSCs (at 80% confluency) were dissociated into single cells by treatment with Accutase (Sigma-Aldrich) at 37°C for 5 mins. The cells were then centrifuged at 100 x g for 3 mins and resuspended in 1ml of STEM Diff Neural induction medium (NIM) (Stem cell technologies, Catalog # 05835), supplemented with 10 μM ROCK inhibitor (Selleckchem, Cat # S1049). The cells were counted using a hemocytometer, and 3 x 10^6^ cells were suspended in 1 ml of NIM medium, supplemented with 10 μM ROCK inhibitor. The resuspended cells were carefully added into our custom-made 5D microwell plate containing 1 ml of NIM, reaching a final volume of 2 ml in each well of the microwell plate. The cell suspension was carefully mixed by pipetting 5-6 times to distribute the cells evenly. After sealing the plate with parafilm, the plate was centrifuged at 1000 rpm for 8 minutes. The plate was then turned and centrifuged again at 1000 rpm for 8 minutes to ensure the cell seeding at the center of the microwells, assuming each microwell receives approximately 10,000 cells. The parafilm was removed, and the plate was incubated at 37°C overnight.

#### Neurosphere formation and transfer to spinner flasks

For the next five days, partial media change was done by replenishing 0.8 ml of NIM with 1 ml of NIM daily without the ROCK inhibitor. During this period, the iPSCs formed strong interconnections, which gave them a 3D appearance known as neurospheres. After 5 days of partial medium change, the neurospheres were detached from the microwells by pipetting them slowly with DMEM/F12 medium (Gibco). Next, these neurospheres were transferred with a cut 1000 µl pipette tip into a 40μm cell strainer (Corning, Cat

\# CLS431750). This procedure was repeated until all the neurospheres were detached and carefully transferred. The cell strainer was then inverted, and the neurospheres were released into a 10 cm petri-dish using approximately 5-10 ml of the Neurosphere Medium consisting of 1:1 DMEM/F12 and Neural Basal medium, supplemented with N2 (1:200), B27 w/o Vitamin A (1:100) (Thermo-scientific) 0.05mM MEM non-essential amino acids (Gibco), L-glutamine (1:100, Gibco), Pen-strep (100 μg/ml each), Insulin (1.6g/L, Sigma Aldrich), 0.05 mM β-Mercaptoethanol (Life Technologies).. Once all the neurospheres were released, they were transferred into 500 ml spinner flasks (Integra biosciences, Cat # 10610762) containing 75 ml of the Neurosphere organoid medium. The flasks were then incubated at 37°C on the magnetic platform. The neurospheres were subjected to a rotation setting at a constant speed of 25 rpm. Half media change was performed once every three days. After 4 days, the Neurosphere medium was switched to the brain organoid differentiation medium. This time point was marked as ‘day 0’.

After Day 20, the organoids were further cultured in the Human brain organoid maturation medium devoid of the SMAD inhibitors (SB431542 and Dorsomorphin). The brain organoids were allowed to mature and analyzed at different time points for maturation markers. Table 1 below lists media composition and incubation duration.

### Time-resolved scRNA-seq of Hi-Q organoids

For scRNA library construction, we used the Chromium™ Single Cell 3’ Reagent Kits v3. Single nucleus suspensions in 1XPBS containing 0,04% BSA (700-1200 nuclei/µL concentration) have been checked for viability, debris, and cell aggregates. To achieve single-cell resolution, the cells are delivered at a limiting dilution, such that the majority (∼90-99%) of generated GEMs contain no cell. In contrast, the remainder essentially includes a single cell. Because of the complex composition of organoids, we aim to target 2.000-3.000 cells per sample. Upon dissolution of the Single Cell 3’ Gel Bead in a GEM, primers containing (i) an Illumina R1 sequence (read 1 sequencing primer), (ii) a 16 bp 10x Barcode, (iii) a 12 bp Unique Molecular Identifier (UMI) and (iv) a poly-dT primer sequence are released and mixed with cell lysate and Master Mix. Incubation of the GEMs then produces barcoded, full-length cDNA from poly-adenylated mRNA. After incubation, the GEMs are broken, and the pooled fractions are recovered. Silane magnetic beads remove excess biochemical reagents and primers from the post-GEM reaction mixture. Full-length, barcoded cDNA is then amplified by PCR to generate sufficient mass for library construction.

Enzymatic Fragmentation and Size Selection optimize the cDNA amplicon size before library construction. R1 (read 1 primer sequence) is added to the molecules during GEM incubation. P5, P7, a sample index, and R2 (read 2 primer sequence) are added during library construction via End Repair, A-tailing, Adaptor Ligation, and PCR. The final libraries contain the P5 and P7 primers used in Illumina bridge amplification. A Single Cell 3’ Library comprises standard Illumina paired-end constructs that begin and end with P5 and P7. We allocated Illumina NovaSeq6000 flowcells to sequence with the first read 28nt (cell-specific barcode and UMI) and generate with the second read 90nt 3’mRNA transcriptome data. Using the v3 chemistry version, 25kreads/nucleus are sufficient for comprehensive scRNA analysis. Single-cell RNA-seq of five batches of organoids was performed using the 10× Chromium pipeline.

Reads were mapped to the human genome (GRCh38), and count matrices were generated using CellRanger. Python notebooks were used to analyze the count matrices by custom functions and the package Scanpy v1.8.2 (Wolf, Angerer and Theis, 2018). Raw single-cell count matrices have been produced for each batch of organoids at the three time points. Batches of the same time point have been concatenated, and possible doublets have been filtered out using the scrublet package (Wolock, Lopez, and Klein, 2019). The three matrices have been then concatenated and filtered. Cells were kept if expressing between 200 and 5000 genes, having between 1000 and 40000 counts, and showing lower percent mitochondrial counts than 8%. Counts have been normalized and log-transformed.

Then, the set of highest variable genes has been detected. The total number of counts per cell and the percentage of mitochondrial counts have been regressed in the data. The PCA dimensions have been calculated, and the harmony package has smoothed the batch effect between time points ^43^. Additionally, the *k*-nearest neighbor was computed by the bbknn algorithm ^44^. The 2-dimension representation of such a network has been calculated as a force-directed graph by the Force Atlas2 algorithm ^45^. A Lasso regression model has been trained on cell type labels of a published dataset of embryonic human brain (Nowakowski *et al.*, 2017). Such a model has been used to score labels on each cell in the Hi-Q dataset. We assigned to each cell the label with the higher prediction score obtained. Such cell-type labels have been used to compare batches between the same and different time points. The proportions of each cell type label served as features for principal coordinate analysis to determine distances between samples. The Leiden algorithm and marker genes have calculated unbiased clusters by t-test. The pseudotime ordering was computed using the diffusion pseudotime (DPT) algorithm by setting the proliferative radial glia cells as roots ^46^.

### Hi-Q brain organoid fixation, permeabilization, antigen retrieval, and blocking

Whole brain organoids were fixed for 1 hr using 4% Paraformaldehyde at 37°C. The organoids were then washed with 1x PBS containing 30mM glycine (2-3 times), permeabilized with 1x PBS glycine containing 0.5% Triton X 100, 0.1% Tween 20 for 15 mins, and further blocked with 1x PBS glycine containing 0.5% Fish Gelatin for 1 hour at room temperature (RT). The organoids were treated with 1% SDS for 5 mins for nuclear staining and washed 3x times with 1x PBS glycine. The organoids were then blocked, as described previously.

Brain organoids were first fixed with 4% paraformaldehyde for 20 mins for organoid tissue sections and then suspended in 30% sucrose overnight at 4°C, allowing them to sink. The following day, the organoids were embedded in a tissue-embedding solution containing (7.5% gelatin and 10% sucrose in PBS) ^47^. Next, the embedded organoids were placed on dry ice for rapid freezing and transferred to −80°c overnight. Finally, the frozen organoids were subjected to cryosectioning to obtain 20µm slices using Cryostat Leica CM3050 S.

### Immunofluorescent staining

The Hi-Q organoids (whole mounts or tissue sections) were fixed and blocked as described above. The organoids were then moved to 2.0 ml Eppendorf tubes for immunostaining for whole mounts. Mouse anti-Nestin (1:100, Novus Biologicals), mouse anti-SOX2 (1:50, Abcam), rabbit anti-Arl13b (1:100, Proteintech), rabbit anti-Doublecortin (1:100, Synaptic Systems), rabbit anti-MAP2 (1:200, Proteintech), rabbit anti-synapsin-1 (1:200, Cell Signalling),Rabbit anti-TUJ1(1:400, Sigma - Aldrich), Mouse anti-acetylated tubulin (1:400, Sigma-Aldrich), Mouse anti-Tau (1:100, DSHB), Rat anti-CTIP2 (1:300, Abcam), Rabbit anti-PCP4 (1:100, Proteintech), Rabbit anti-pH2AX (1:400, Cell signalling), Mouse anti-Actin (1:100, R&D systems), rabbit anti-PSD95 (1:100, Proteintech), mouse anti-PAX6 (1:100, Proteintech), mouse anti-p-vimentin (1:200, Abcam) and TUNEL staining kit (Thermo Fischer). We used Alexa Fluor Dyes conjugated with goat/donkey anti-mouse, anti-rabbit, or anti-rat (1:1000, molecular probes, Thermo Fisher, USA) for secondary antibodies. In addition, DAPI 1 μg/ml (Sigma Aldrich, USA) was used to stain the nucleus.

## Whole organoid tissue clearing and mounting

Hi-Q organoids were tissue-cleared after immunostaining using increasing concentrations of ethanol (50%, 70%, and 100%) (PMID: 32521263). The organoids were treated with different percentages of ethanol for approximately 5-6 minutes at RT with constant mixing. After treatment with 100% ethanol, organoids were removed and allowed to dry till all the ethanol evaporated. Next, the organoids were cleared by adding 100 μl of ethyl cinnamate (Sigma-Aldrich, Cat. 8.00238) for 10 minutes. Once cleared, the organoids were transferred with a cut 1000 µl pipette tip to the μ-slide angiogenesis chamber (ibid, Cat # 81506), which could readily be imaged.

### Confocal microscopy

Tissue sections or whole-mount tissue-cleared organoids were imaged using a Zeiss LSM 880 confocal microscope. Our experiments used 405, 488, 561, or 633 nm laser lines and were imaged using objective 10x/0.3, 20x/0.8 M27, and 40x/1.3 oil objectives. After tissue clearing, the whole-brain organoids were suspended in ethyl cinnamate and imaged in the μ-slide angiogenesis chamber. Z-series were obtained for both whole organoids and organoid slices, and the slice interval between the stacks was approximately 1-2 µm. The raw images acquired were processed using ImageJ, Adobe Photoshop 2020, and Adobe Illustrator 2020. Movies of organoid Z-stack images were processed using ImageJ.

### Statistical analysis

The statistical analyses were performed using GraphPad Prism (version 9). Most experiments were carried out in triplicates and analyzed using non-parametric one-way ANOVA or Student’s t-test. The Post hoc test included primarily Tukey’s Test. All values were expressed as mean± sem. ‘N’ represents the number of organoids or components, and ‘n’ represents the number of experimental replicates. Statistical significance is represented as ns-nonsignificant, *p<0.1, **p<0.01, ***p<0.001.

### Calcium Imaging

Brain organoids were cut into two halves, put with the cut surface down onto an experimental chamber, and fixed with a grid. Preparations were perfused with artificial cerebrospinal fluid (ACSF) at room temperature (20-22°C), containing (in mM): 125 NaCl, 2.5 KCl, 2 CaCl2, 1 MgCl2, 1.25 NaH2PO4, 26 NaHCO3, and 20 glucose, bubbled with 95% O2 and 5% CO2, resulting in a pH 7.4 and an osmolarity of 310 ± 5 mOsm/l (all chemicals purchased from Sigma-Aldrich (Munich, Germany), except for indicator dyes and tetrodotoxin (TTX; BioTrend Chemicals AG, Zürich, Switzerland)). Experiments were performed at room temperature as well. For imaging of intracellular Ca2+, the membrane-permeable form of the chemical Ca2+ indicator dye Oregon Green BAPTA 1 (OGB1-AM; Thermo-Fisher Scientific, Waltham, USA) was dissolved in HEPES-buffered saline composed of (in mM) 125 NaCl, 2.5 KCl, 2 CaCl2, 2 MgCl2, 1.25 NaH2PO4 and 25 HEPES, pH 7.4 (adjusted with NaOH) OGB1-AM was then bolus-injected (5 s/2-3 PSI) into organoid preparations at a depth of about 30-50 µm, this injection was repeated at up to 10 adjacent positions in the field of view. Imaging experiments were commenced 30 min after dye injection, allowing for de-esterification of OGB1-AM.

Wide-field Ca2+ imaging was performed using a digital variable scan system (Nikon NIS-Elements v4.3, Nikon GmbH Europe, Düsseldorf, Germany). It was equipped with a 40x/N.A. 0.8 LUMPlanFI water immersion objective (Olympus Deutschland GmbH, Hamburg, Germany) and an orca FLASH V2 camera (Hamamatsu Photonics Deutschland GmbH, Herrsching, Germany). OGB1 was excited at 488 nm, and emission was collected >500 nm at 14-20 Hz. Fluorescence emission was recorded from defined regions of interest (ROI) representing cell bodies employing NIS-Elements software (Nikon GmbH Europe, Düsseldorf, Germany). For background correction, an ROI in the field of view apparently devoid of cellular structures was chosen, and its fluorescence emission was subtracted from that derived from each cellular ROI. Background-corrected signals were analyzed offline using OriginPro Software (OriginLab Corporation v9.0, Northampton, USA). Changes in intracellular Ca2+ were expressed as ΔF/F0, representing changes in fluorescence over time (ΔF) divided by the average baseline of each ROI (F0). Only events with peak amplitudes higher than two times the standard deviation of the baseline noise were analyzed. Glutamate or GABA was applied by perfusing preparations with ACSF, to which the neurotransmitters were added at a concentration of

1 mM. Pharmacological antagonists were dissolved in aCSF and washed 15 min before the experiment, allowing sufficient blocking of the target receptor or channel.

After wide-field Ca2+ imaging, some preparations were transferred to an upright confocal laser scanning microscope (Nikon C1 Eclipse E600FN, Nikon GmbH Europe, Duesseldorf, Germany) with a 40x/N.A. 0.8 LUMPlanFI water immersion objective (Nikon GmbH Europe, Düsseldorf, Germany) and a 488 nm argon laser (Melles Griot GmbH, Bensheim, Germany). Z-stacks of the imaged region were generated (up to 240 images at a step size of 0.1 µm). They were post-processed, employing deconvolution software to delineate cellular morphology (Huygens Professional, SVI imaging, Hilversum, Netherlands).

Unless otherwise specified, data are presented in Tukey box-and-whisker plots indicating median (line), mean (square), interquartile range (IQR; box), and standard deviation (whiskers). In addition, all individual data points are shown in grey underneath the Tukey plots. Data were statistically analyzed by one-way ANOVA followed by post hoc Bonferroni test. The following symbols are used to illustrate the results of statistical tests in the figures: *: 0.01 ≤ p < 0.05; **: 0.001 ≤ p < 0.01; ***: p < 0.001. “n” represents the number of cells analyzed; “N” represents the number of individual experiments/brain organoids. Each series of experiments was performed on at least three different organoids.

### Data Analysis and Statistics (related to calcium imaging experiments)

Unless otherwise specified, data are presented in Tukey box-and-whisker plots indicating median (line), mean (square), interquartile range (IQR; box), and standard deviation (whiskers). In addition, all individual data points are shown in grey underneath the Tukey plots. Data were statistically analyzed by one-way ANOVA followed by post hoc Bonferroni test. The following symbols are used to illustrate the results of statistical tests in the figures: *: 0.01 ≤ p < 0.05; **: 0.001 ≤ p < 0.01; ***: p

< 0.001. “n” represents the number of cells analyzed; “N” represents the number of individual experiments/brain organoids. Each series of experiments was performed on at least three different organoids.

### Cryopreservation, thawing, and re-culturing of Hi-Q brain organoids

The required number of immature brain organoids on day 8 was carefully aspirated from the spinner flasks and placed in a 6 cm petri dish containing a 5 ml organoid medium. With the help of a cut 1000 µm filter tip, these brain organoids were transferred into freezing vials containing commercial freezing media (CS10, Stemcell Technologies). Approximately 10 brain organoids were frozen per vial containing 1 ml of CS10. The freezing mixture containing the organoids was kept on ice for 5 minutes before transferring them in an isopropanol chamber at −80°c for controlled freezing. 24 to 48 hours later, these frozen vials were transferred to liquid nitrogen tanks. The frozen organoids were re-thawed after 5-7 days to check for organoid viability, morphology, and functionality. The frozen organoids were rapidly thawed at 37°C and then transferred with a cut 1ml sterile pipette tip into a 6 cm dish containing 5 ml organoid medium. These organoids were placed at 37°C in static conditions for 48 hours to recover from thawing. On Day 3, the organoids were carefully transferred to a spinner flask containing 75 ml of brain organoid medium and allowed to mature in the stirred conditions described above.

### Glioblastoma invasion assays and drug screening in Hi-Q organoids

Glioblastoma primary cell line 450-cherry was maintained as described previously ^23,24^. Invasion assays were performed by single-cell inoculation or by fusion of GSC spheres with our brain organoids. Plate and liquid handling were performed using a high-throughput screening workstation for organoid screening. Three plates of the Target Selective Screening Library (Selleckchem) were used for the screening (180 validated selective inhibitors dissolved in DMSO, 1 mM stock solution). Hi-Q organoids (untagged hiPSC cells with GBM-Cherry) were cultivated in CellCarrier Spheroid ULA 96-well Microplates (PerkinElmer). 72 h after organoids were placed in the plates, they were either treated with a 5 µM compound (dissolved in DMSO) or alone. In parallel, one plate was treated with control compounds. The final DMSO volume concentration was kept below 0.5%. Organoids were cultivated at 37°C with 5% CO2 and imaged after 24 h and 72 h. Imaging of two channels (Brightfield and DsRed) was performed with the automated Operetta® High-Content microscope (Perkin Elmer)—2x objective, 1 field per well, 20 planes.

Image analysis was performed using the Columbus software (PerkinElmer). The following analysis steps in Columbus are described: as an input image, a maximum projection of the different planes was used. Brightfield signal was inverted via the ‘Filter image’ tool and then used to detect the organoid (Find Nuclei, Method: B, Common Threshold: 0.4, Area: > 5000 µm², Splitting Coefficient: 7, Individual Threshold: 0.4, Contrast: > 0.1). The DsRed channel was smoothened using ‘Mean Filter’ and then used to detect invasion of glioma stem cells via the ‘Find spot’ tool. (Method: A, Relative Spot Intensity: > 0.08, Splitting Sensitivity: 0.9, Calculate Spot Properties). Compounds were defined as hit compounds if the compound showed only 0 or 1 spot in both time points, *i.e.,* 24 h and 72 h.

---

**Figure S1.**
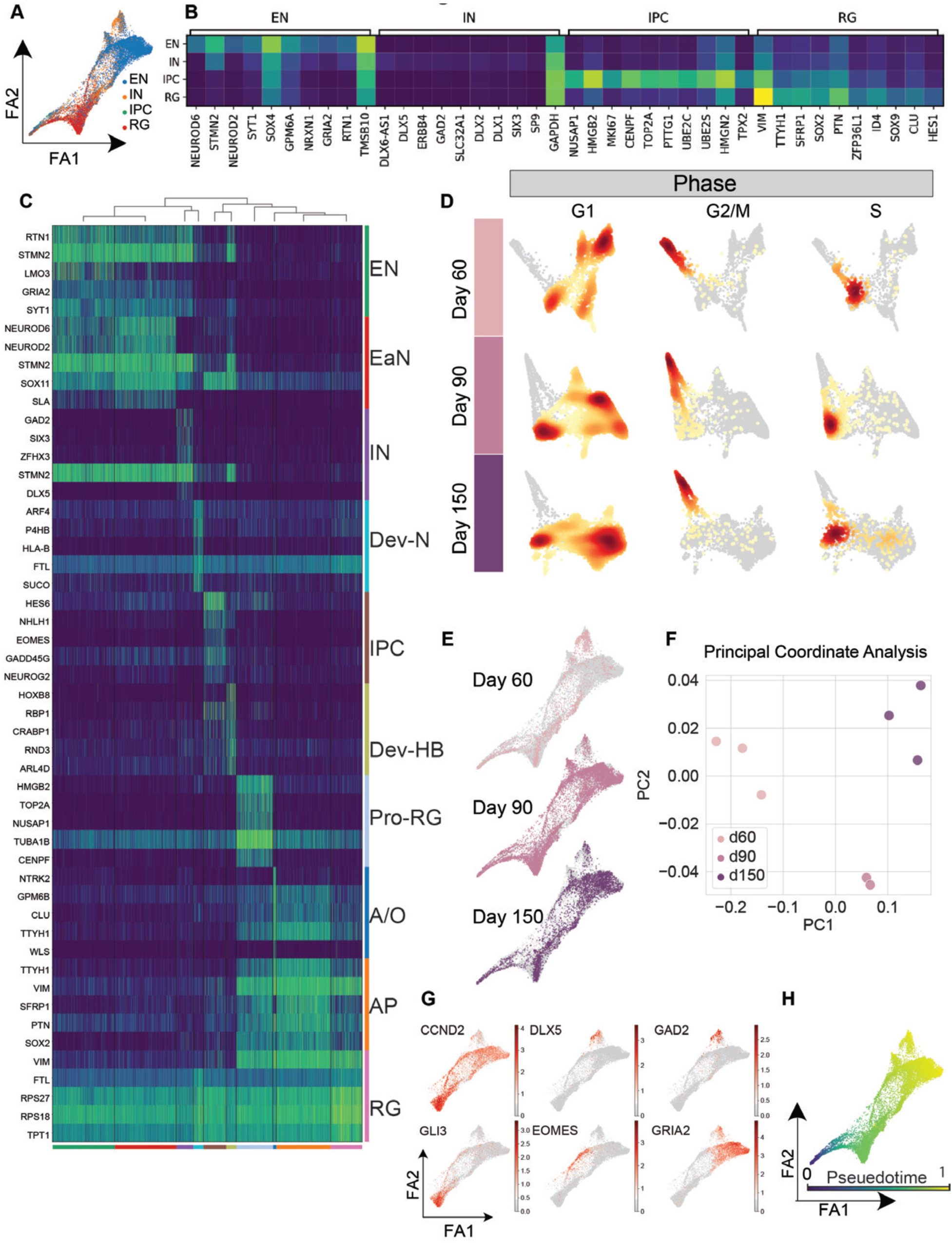
Cell type heterogeneity in Hi-Q brain organoids across different time points of maturation (Related to Figure 2). **A.** Force Altas (FA) 2D representation of the neighbor graph of Hi-Q brain organoids from day 60, day 90, and day 150 datasets with cell type predicted labels. **B.** Matrixplot showing the average gene expression of the marker genes (top 10) of each cell type label in the integrated dataset. Markers are calculated by pairwise Wilcoxon rank sum test of each group against the rest of the cells. **C.** Heatmap showing the expression of marker genes calculated on unbiased Leiden clustering. **D.** FA embedding of day 60, day 90, and day 150 separately indicates the density of cells expressing markers characteristic of cell-cycle stages. **E.** FA embedding of the integrated datasets highlighting cells from different collection time points. **F.** Scatterplot reporting each dataset in the function of the first two principal coordinates (PC) calculated from the proportion of cell types predicted in each batch of Hi-Q brain organoids. **G.** FA embedding of the integrated dataset subset. Cells are highlighted by the expression of ventral telencephalon development markers (CCND2, DLX5, and GAD2) and dorsal telencephalon development markers (GLI3, EOMES, and GRIA2). **(H)** FA embedding of the whole integrated dataset highlighted by diffusion pseudotime score computed setting the Pro-RG cluster as root.

**Figure S2:**
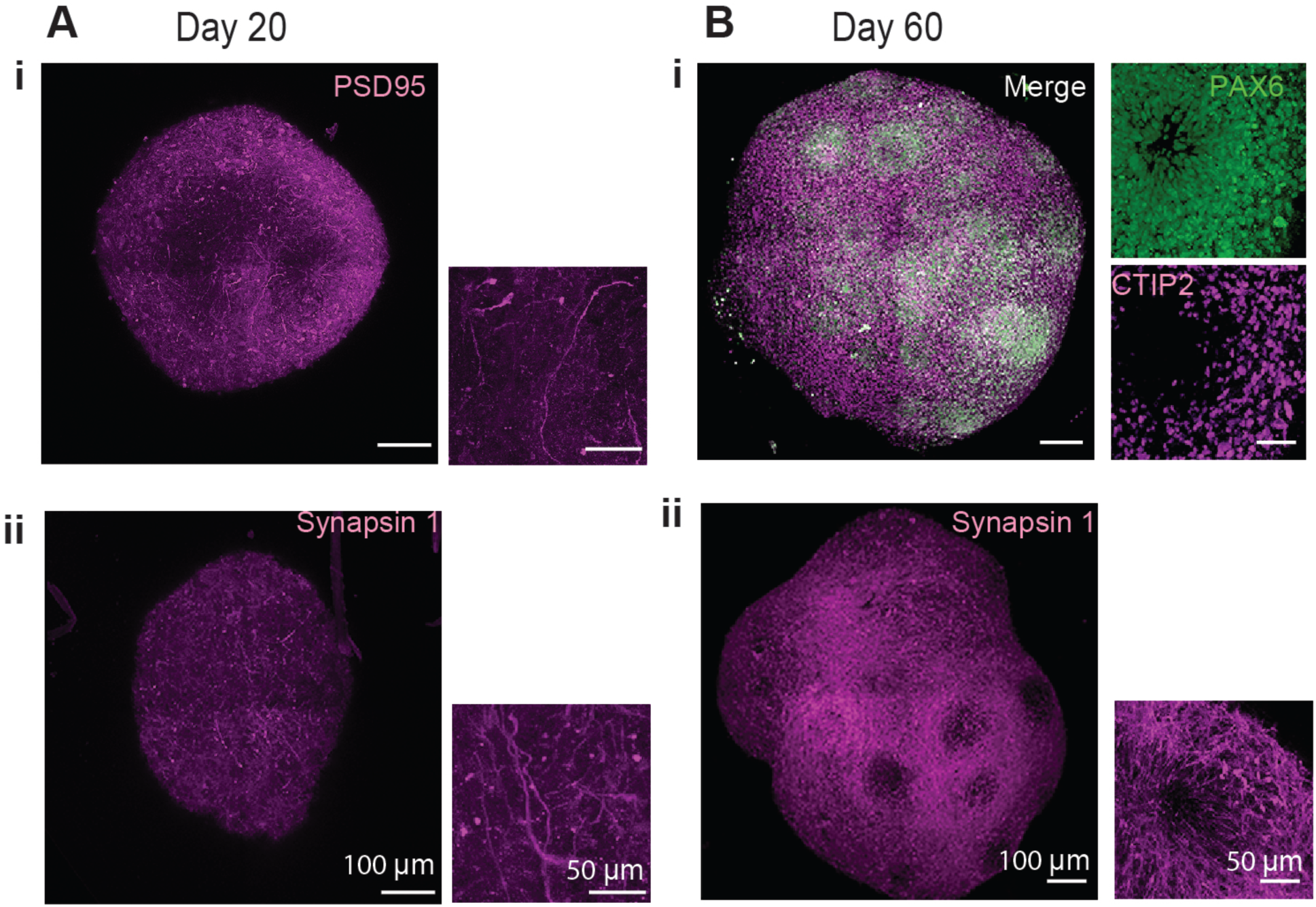
Hi-Q brain organoids mature over time (Related to Figure 3) **A-B.** Tissue clearing and wholemount staining of day 20 **(Ai-ii)** and 60-day **(Bi-ii)** old Hi-Q brain organoids show the presence of PSD95 (magenta), CTIP2 (magenta) and Synapsin 1(magenta). Representative images are shown, and panels show scale bars.

**Figure S3:**
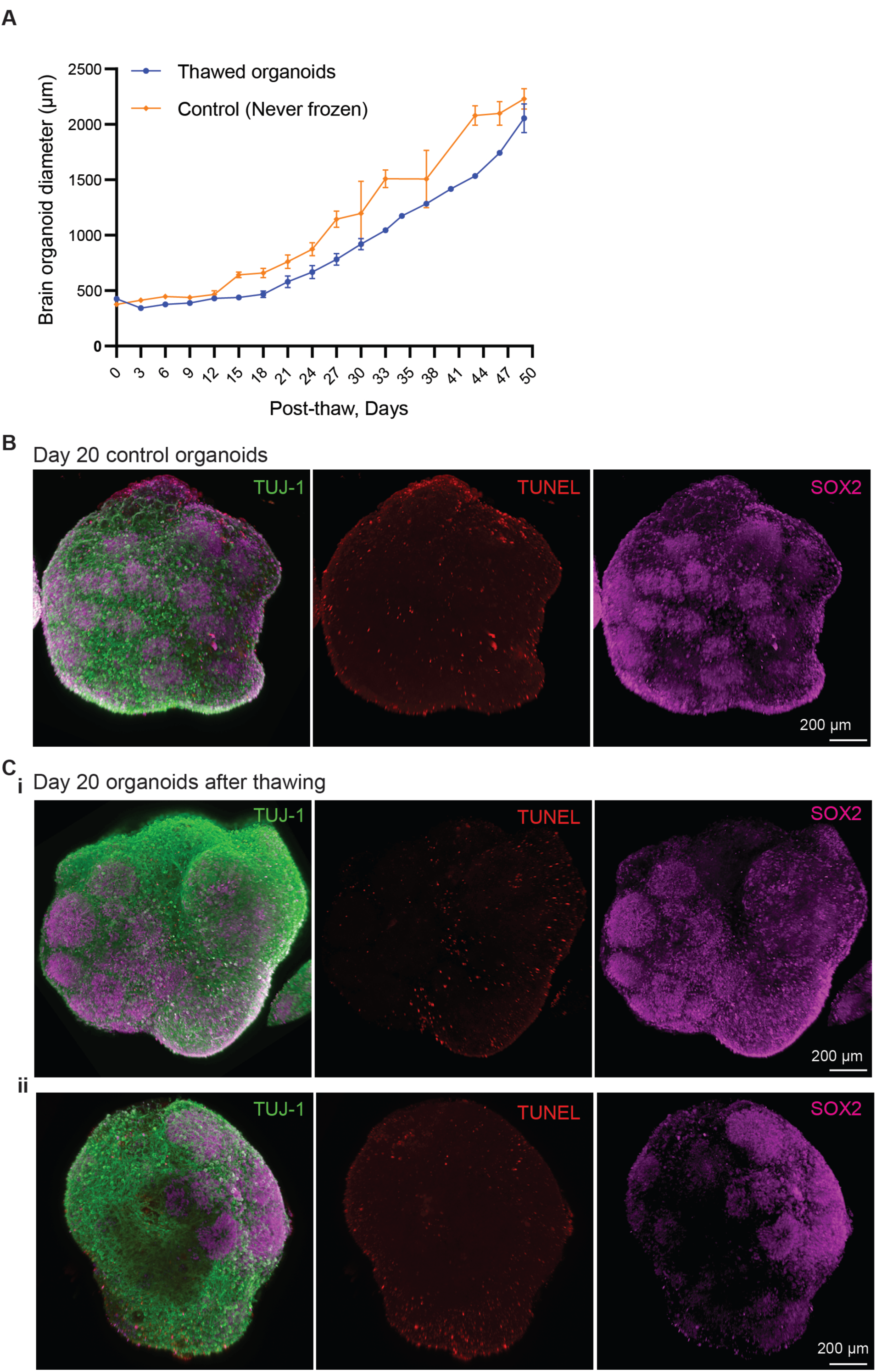
Cryopreservation, thawing, and re-culturing of Hi-Q brain organoids (Related to Figure 5) **A.** Growth kinetics of thawed Hi-Q brain organoids (blue line) compared to control organoids (orange line) that have never been frozen. Each time point shows the average diameter of at least four organoids. **B-C.** Comparison of cytoarchitecture of day-20 Hi-Q brain organoid **(B)** (control, never frozen) and thawed organoids **(C).** Ventricular zone (VZ) is marked by SOX2 (magenta). Neurons are marked by marked by TUJ-1. TUNEL (red) labels dead cells.

**Figure S4:**
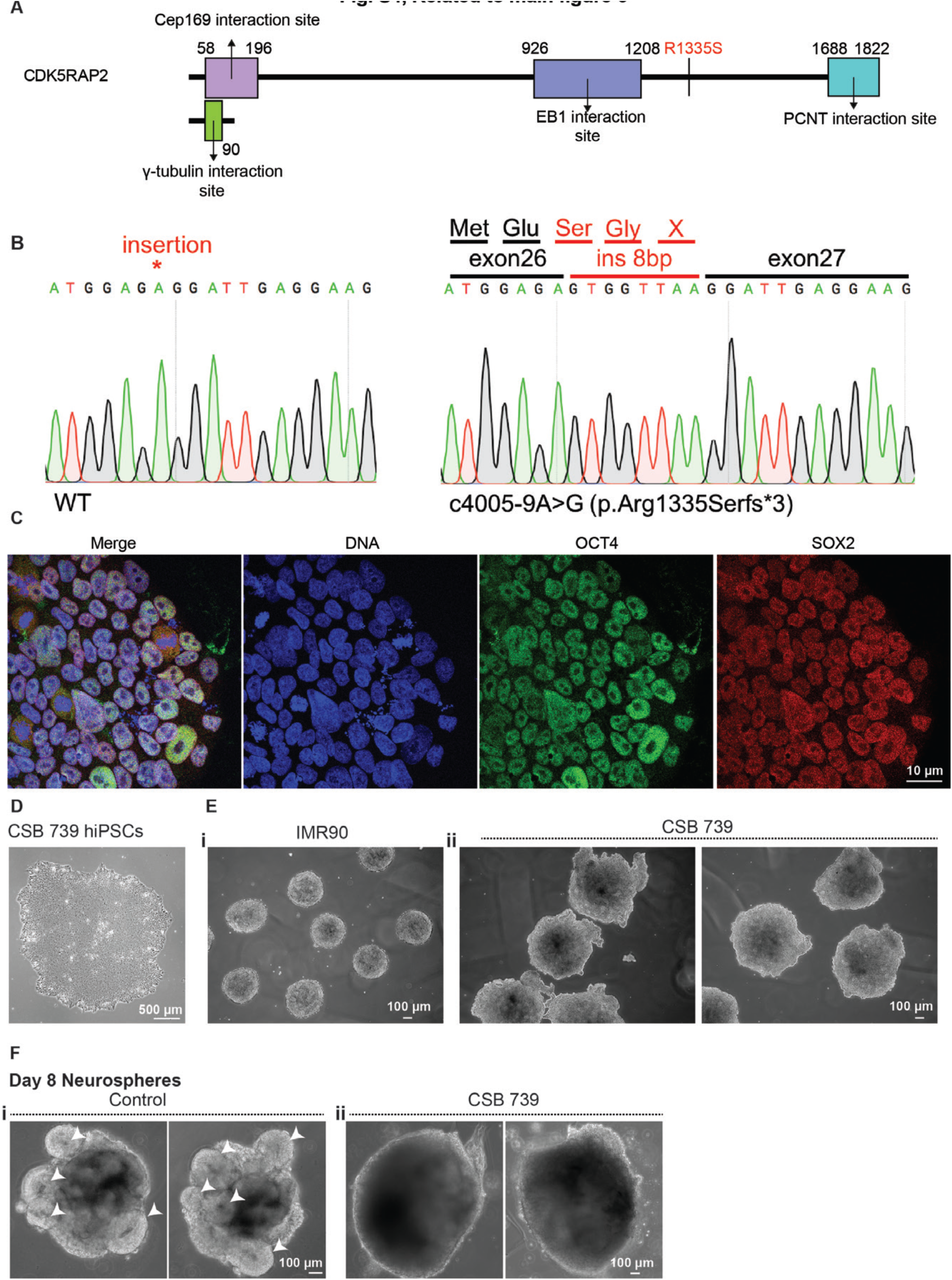
Hi-Q brain organoids model microcephaly caused by a CDK5RAP2 mutation and brain organization defects caused by a CSB mutation (Related to Figure 6) **A.** Schematics of the CDK5RAP2 secondary structure showing protein interaction sites and the frameshift mutation R1335S. **B.** DNA sequence of mutant CDK5RAP2 from patient-derived iPSCs (right) compared to healthy control (left). Nucleotide insertions between exon 26 and 27 are highlighted (red), leading to a frameshift with an amino acid change from R to S. **C.** Patient-derived iPSCs displaying pluripotent markers of OCT4 and SOX2. **D-F.** Comparison of neurospheres derived from healthy control (IMR90) and Cockayne Syndrome (CSB 739; due to a mutation is CSB). Bright field images. Arrowheads in panel F point to well-defined neurospheres in healthy controls. Such structures are not observed in CSB739. Panels show the scale bar.

**Figure S5.**
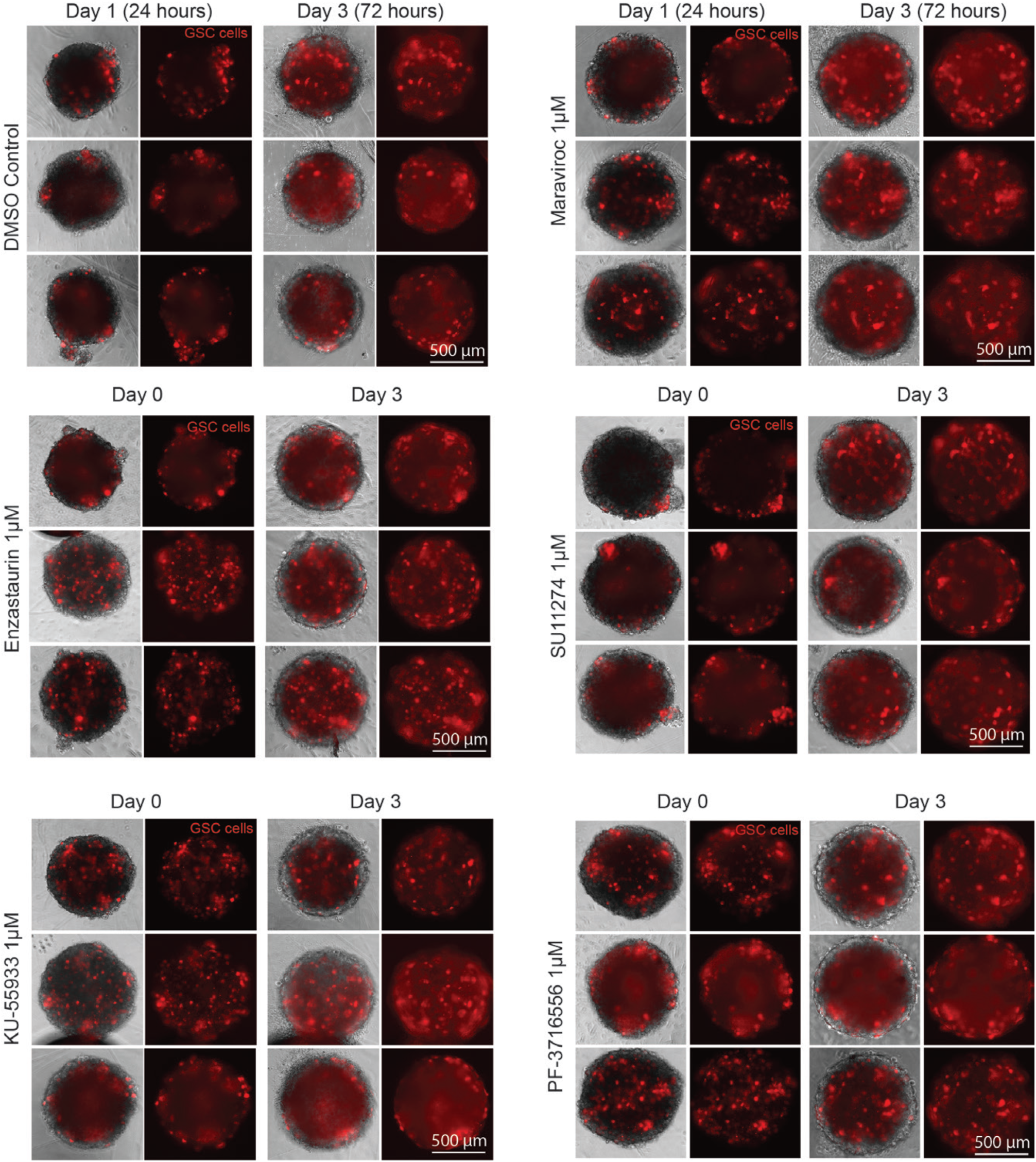
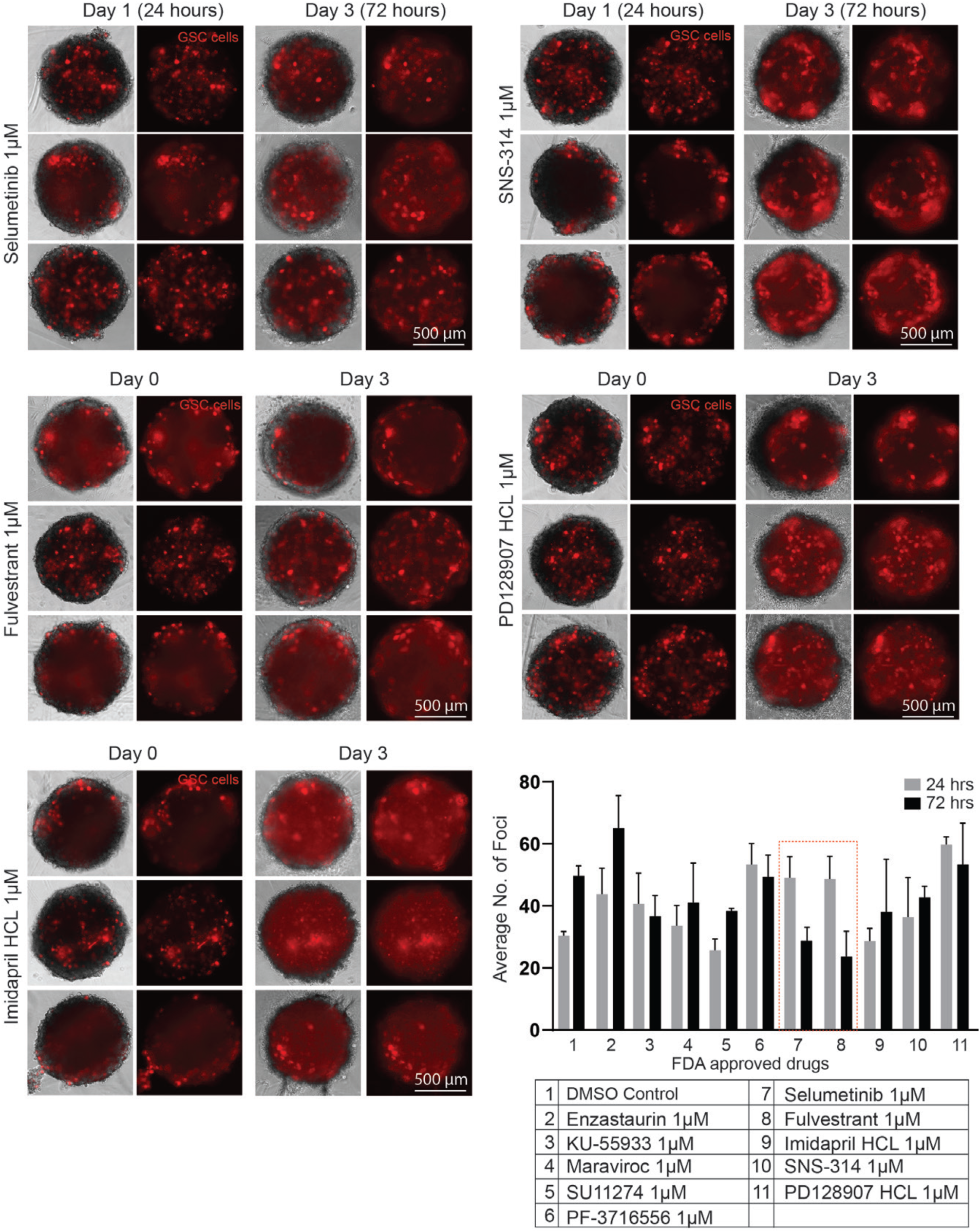
(Part 1 and 2): Secondary screening of compounds that perturb GSCs invasion in Hi-Q brain organoids. Testing of ten compounds (listed in the table at the end of the second part of the figure) for their ability to impair GSCs invasions in day-40 Hi-Q brain organoids. Patient-derived GSCs are labeled with mCherry. All experimental point includes three independent invasion assays with 1 µM of the selected compound. Invasions were measured at two different time points (24 and 72 hours). Note that the time point day 1 (or 24 hours) is 24 hours after seeding GSCs to brain organoids. The GSCs foci were calculated by automated imaging. The bar graph at the bottom quantifies the GSCs foci at 24 and 72 hours. The secondary screen identifies compounds 7 and 8 (Selumetinib and Fulvestrant) inhibiting GSC invasion (The dotted red box).

**Figure S6:**
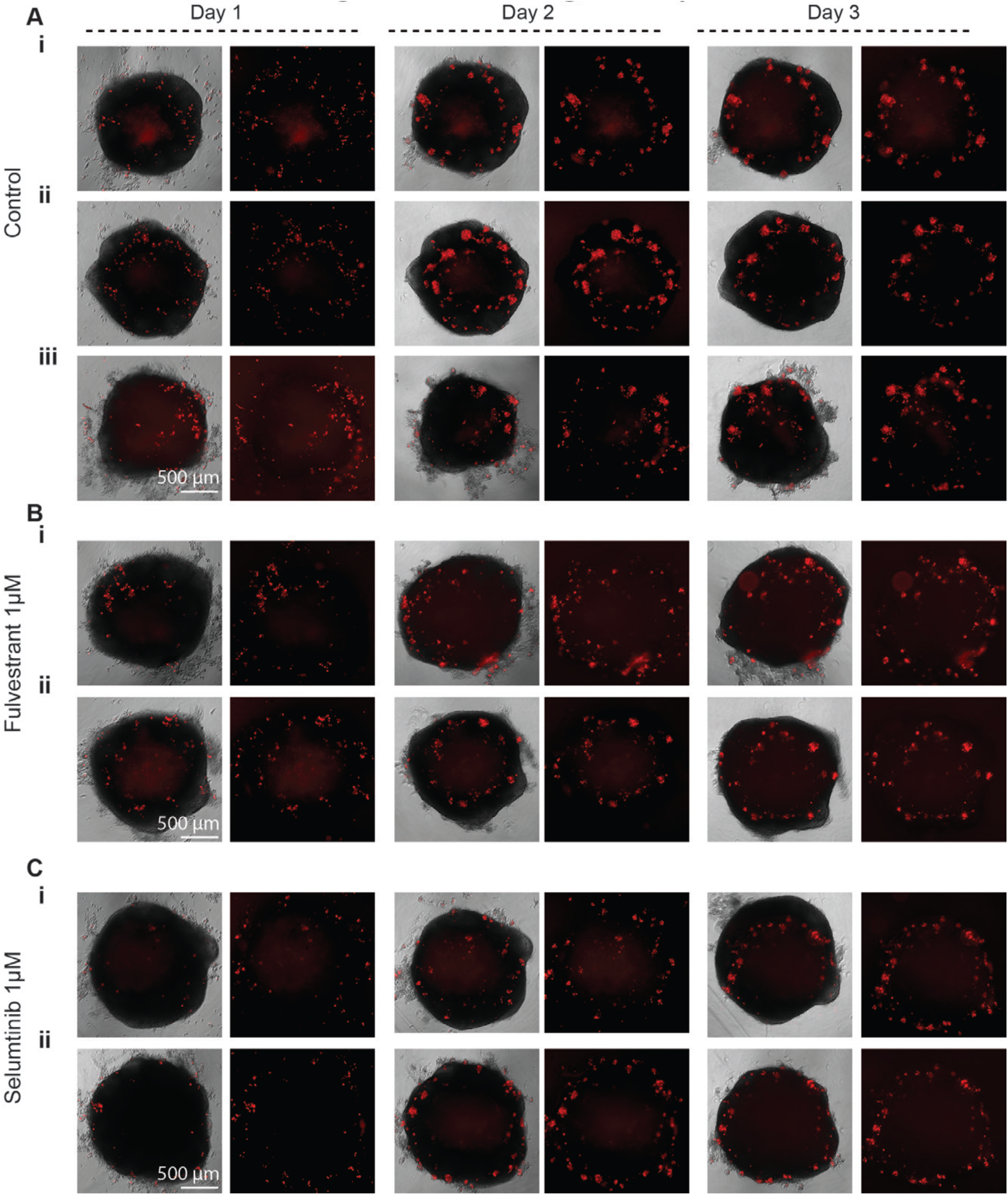
Testing Selumetinib and Fulvestrant for their ability to perturb GSCs invasions in Hi-Q brain organoids (as cell suspension at low resolution) **A.** Kinetics of GSCs (red) invasions at day 0, day 1, and 2. Experiments were conducted in triplicate **(i-iii).** Panels show scale bars. **B-C.** Exposure to 1µM Selumetinib **(i-ii)** and Fulvestrant **(i-ii)** appears to prevent GSC invasions into the organoids. Compared to untreated controls, drug-treated organoids show GSCs at the periphery of organoids. At least two representative organoid images are given. Panels show scale bar.

**Figure S7:**
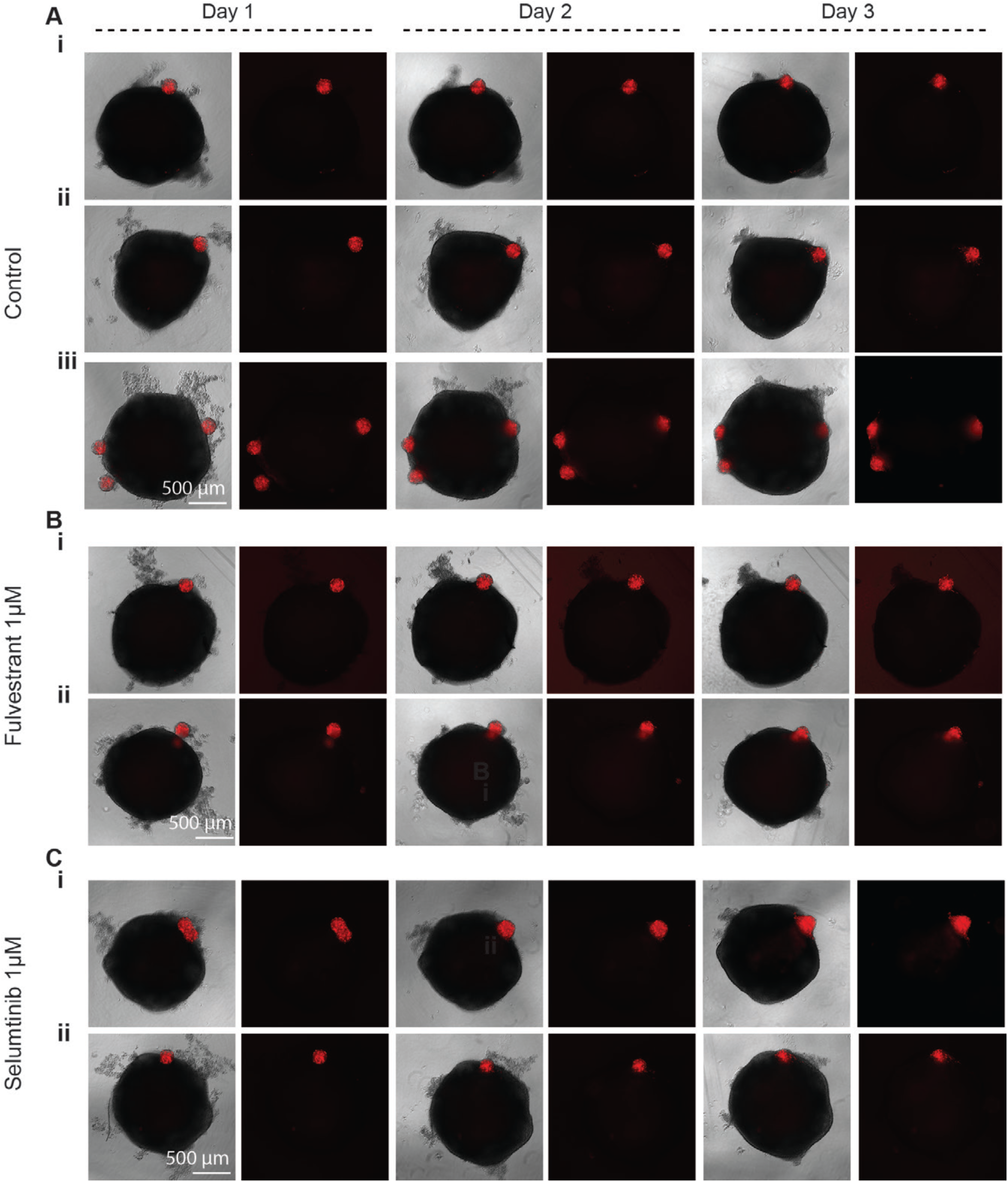
Testing Selumetinib and Fulvestrant for their ability to perturb GSCs invasions in Hi-Q brain organoids (as spheres at low resolution) **A.** Kinetics of GSCs spheres (red) invasions on day 0, day 1, and 2. Experiments were conducted in triplicate **(i-iii).** GSC spheres integrate into organoids. Panels show scale bars. **B-C.** Exposure to 1µM Selumetinib **(i-ii)** and Fulvestrant **(i-ii)** appears to prevent GSC spheres integration to organoids. Compared to untreated controls, drug-treated GSC spheres mainly stay at the edges of organoids. At least two representative organoid images are given. Panels show scale bar.

**Figure S8:**
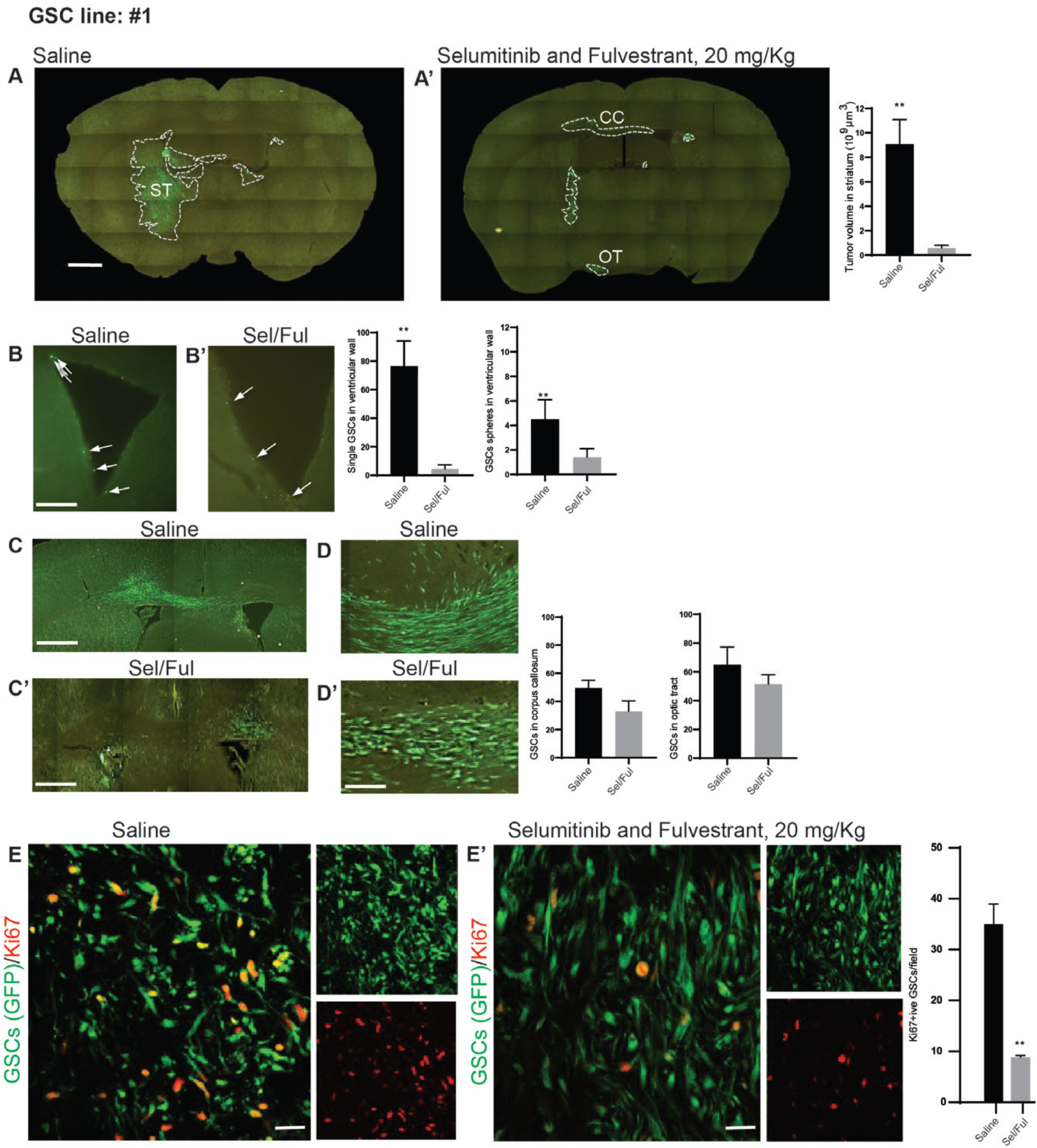
Combined treatment of Selumetinib and Fulvestrant perturb GSC (GSC line #1) invasion in mouse xenografts. **A-A’.** In contrast to saline treatment, 20 mg/Kg treatment of Selumetinib and Fulvestrant **(A’, right)** prevents integration of grafted GSC into the corpus callosum (CC) and optic tract (OT). Drug treatment reduces the tumor volume in the striatum. The bar diagram at right quantifies the tumor volume. Unpaired t-test and Mann-Whitney U test were used. **B-B’.** Compared to saline treatment **(B)**, Selumetinib and Fulvestrant **(B’, right)** prevent integration of grafted GSC into the ventricular wall. The bar diagram at right quantifies the integrated GSCs. Unpaired t-test and Mann-Whitney U test were used. **C-C’.** In contrast to saline treatment **(C)**, Selumetinib and Fulvestrant **(C’, bottom)** prevent grafted GSC integration into the corpus callosum ventricular wall. The bar diagram at right quantifies the integrated GSCs. **D-D’.** Selumetinib and Fulvestrant treatment **(D’, bottom)** prevents grafted GSC integration into the optic tract. Panel **D** (top) shows saline control. The bar diagram at right quantifies the number of GSCs integration between control and treated groups. **E-E’. GSC proliferation between saline and drug treatment.** Selumetinib and Fulvestrant treatment **(E’, right)** significantly reduces the GSC (green) proliferation as probed by Ki67 staining (red)grafted GSC in the optic tract. The bar diagram at right quantifies the GSC proliferation between control and treated groups. Unpaired t-test and Mann-Whitney U test were used.

**Figure S9:**
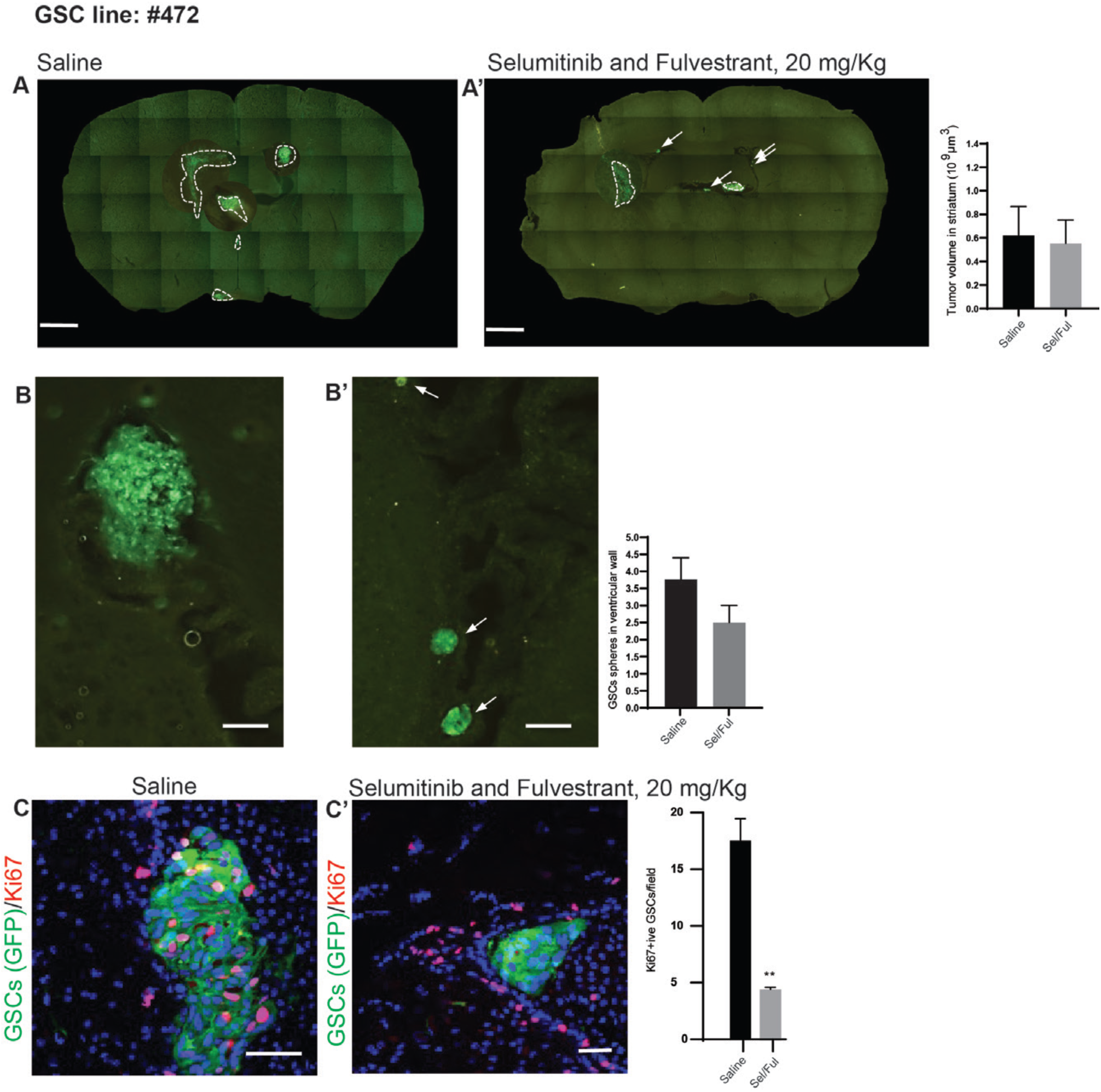
Combined treatment of Selumetinib and Fulvestrant perturb GSC (GSC line #472) invasion in mouse xenografts. **A-A’.** Selumetinib and Fulvestrant-treated animals **(A’, right)** show a reduced tumor volume in the striatum. The saline-treated group is labeled as **A.** The bar diagram at right quantifies the tumor volume. **B-B’.** Compared to saline treatment **(B)**, Selumetinib and Fulvestrant **(B’, right)** prevent the integration of grafted GSC spheres in the ventricular wall. The bar diagram at right shows the quantification. **C-C’. Compared to saline-treated animal groups (C), drug-treated animals (C’) show a reduced proliferation of grafted GSCs proliferation** as probed by Ki67 staining (magenta) grafted GSC in the optic tract. The bar diagram at right quantifies the GSC proliferation between control and treated groups. Unpaired t-test and Mann-Whitney U test were used.

## Figure legend (Movie)

**Movie 1:** Bioreactor equipped with spinning arms to culture Hi-Q brain organoids. The movie shows suspending brain organoids.

**Movie 2 (Related to main** figure 3): Day 20 and day 60 Hi-Q brain organoids showing the distribution of various cell identity markers such as Acetylated a-tubulin, PSD97, SOX2, DCX, Synapsin 1, Tau, MAP2, Actin, P-Vimentin, Nestin and PCP4. Z-series stacks were collected after whole-mount staining, tissue clearing, and confocal imaging. Scale bar 500 µM.

**Movie 3 (Related to main** figure 5**): Control (never frozen) and** thawed Hi-Q brain organoid after cryopreservation showing the cytoarchitecture and distribution of dead cells labeled by TUNNEL. Z-series stacks were collected after whole-mount staining, tissue clearing, and confocal imaging. Scale bar 500 µM.

