## Supplementary material for "Reliability of high-quantity human brain organoids for modeling microcephaly, glioma invasion, and drug screening": Table 2

**Table 2. Organoid batches**

| **Exp/ Batch no.** | **Researcher no.** | **hiPSC cell type** | **No. of neurospheres initiated** | **No. of organoids in spinner flasks** | **Status of experiment** | **Remarks** | **Status of Freeze/thaw** |
| --- | --- | --- | --- | --- | --- | --- | --- |
| 1 | Researcher 1 | IMR90 (WiCell Research Institute, RRID:CVCL_C436) | 370 | 350 | Successful | Neurospheres lost during neural induction and transfer to spinner flasks | Organoids were not frozen |
| 2 | Researcher 1 | IMR90 (WiCell Research Institute, RRID:CVCL_C436) | 1080 | 950 | Successful | Fused organoids were discarded | Organoids were not frozen |
| 3 | Researcher 1 | IMR90 (WiCell Research Institute, RRID:CVCL_C436) | 370 | 298 | Successful | - | 20 organoids were freeze-thawed |
| 4 | Researcher 1 | IMR90 (WiCell Research Institute, RRID:CVCL_C436) | 740 | 725 | Successful | Organoids disintegrated in spinner flasks | Organoids were not frozen |
| 5 | Researcher 1 | IMR90 (WiCell Research Institute, RRID:CVCL_C436) | 555 | 520 | Successful | Neurospheres lost during neural induction | Organoids were not frozen |
| 6 | Researcher 1 | IMR90 (WiCell Research Institute, RRID:CVCL_C436) | 555 | 530 | Successful | Fused organoids were discarded from spinner flasks | 20 organoids were freeze-thawed |
| 7 | Researcher 1 | IMR90 (WiCell Research Institute, RRID:CVCL_C436) | 740 | 700 | Successful | Fused organoids were discarded from spinner flasks | Organoids were not frozen |
| 8 | Researcher 1 | Tubulin-GFP tagged hiPSCs (Coriell, Cat# AICS-0012) | 370 | 345 | Successful | Neurospheres lost during neural induction | Organoids were not frozen |
| 9 | Researcher 1 | Tubulin-GFP tagged hiPSCs (Coriell, Cat# AICS-0012) | 185 | 180 | Successful | Fused organoids were discarded | Organoids were not frozen |
| 10 | Researcher 1 | Tubulin-RFP tagged hiPSCs (Coriell Cat# AICS-0031-035) | 370 | 360 | Successful | Fused organoids were discarded | Organoids were not frozen |
| 11 | Researcher 1 | Tubulin-RFP tagged hiPSCs (Coriell Cat# AICS-0031-035) | 185 | 175 | Successful | Neurospheres lost during transfer | Organoids were not frozen |
| 12 | Researcher 1 | Crx-ips (PMID: [30100409](https://pubmed.ncbi.nlm.nih.gov/30100409)) | 370 | 355 | Successful | Neurospheres lost during transfer | Organoids were not frozen |
| 13 | Researcher 1 | Crx-ips (PMID: [30100409](https://pubmed.ncbi.nlm.nih.gov/30100409)) | 185 | 145 | Successful | Organoids disintegrated (Reason unknown) | Organoids were not frozen |
| 14 | Researcher 1 | Crx-ips (PMID: [30100409](https://pubmed.ncbi.nlm.nih.gov/30100409)) | 185 | 170 | Successful | Neurospheres lost during transfer | Organoids were not frozen |
| 15 | Researcher 1 | CDK5RAP2 ipsc | 185 | 180 | Successful | Neurospheres lost during transfer | Organoids were not frozen |
| 16 | Researcher 1 | CDK5RAP2 ipsc | 185 | 160 | Successful | Neurospheres lost during transfer / Fused organoids were discarded | Organoids were not frozen |
| 17 | Researcher 1 | CDK5RAP2 ipsc | 185 | 180 | Successful | - | Organoids were not frozen |
| 18 | Researcher 1 | CSB-GM739 (PMID: [22904069](https://pubmed.ncbi.nlm.nih.gov/22904069)) | 370 | 350 | Successful | Fused organoids were discarded | Organoids were not frozen |
| 19 | Researcher 1 | CSB-GM739 (PMID: [22904069](https://pubmed.ncbi.nlm.nih.gov/22904069)) | 185 | 170 | Successful | Neurospheres lost during transfer | Organoids were not frozen |
| 20 | Researcher 1 | CSB-GM739 (PMID: [22904069](https://pubmed.ncbi.nlm.nih.gov/22904069)) | 185 | 175 | Successful | Fused organoids were discarded | Organoids were not frozen |
| 21 | Researcher 1 | CSB-GM739 (PMID: [22904069](https://pubmed.ncbi.nlm.nih.gov/22904069)) | 185 | 180 | Successful | Fused organoids were discarded | Organoids were not frozen |
| 22 | Researcher 2 | Tubulin-GFP tagged hiPSCs (Coriell, Cat# AICS-0012) | 370 | 325 | Successful | Neurospheres lost during neural induction | Organoids were not frozen |
| 23 | Researcher 2 | Tubulin-GFP tagged hiPSCs (Coriell, Cat# AICS-0012) | 370 | 350 | Successful | Fused organoids were discarded | Organoids were not frozen |
| 24 | Researcher 2 | Tubulin-RFP tagged hiPSCs (Coriell Cat# AICS-0031-035) | 370 | 350 | Successful | Neurospheres lost during neural induction | Organoids were not frozen |
| 26 | Researcher 2 | IMR90 (WiCell Research Institute, RRID:CVCL_C436) | 350 | 100 | Partially successful | Neurospheres lost during neural induction andmost organoids were collapsed | Organoids were not frozen |
| 25 | Researcher 2 | IMR90 (WiCell Research Institute, RRID:CVCL_C436) | 400 | 350 | Successful | Fused organoids were discarded | Organoids were frozen |
| 25 | Researcher 2 | Tubulin-GFP tagged hiPSCs (Coriell, Cat# AICS-0012) | 750 | 700 | Successful | Neurospheres lost during neural induction | Organoids were not frozen |
| 26 | Researcher 3 | IMR90 (WiCell Research Institute, RRID:CVCL_C436) | 500 | 450 | Successful | Neurospheres lost during neural induction | Organoids were not frozen |
| 27 | Researcher 3 | Tubulin-GFP tagged hiPSCs (Coriell, Cat# AICS-0012) | 550 | 500 | Successful | Fused organoids were discarded from spinner flasks | Organoids were freeze-thawed |
| 28 | Researcher 3 | IMR90 (WiCell Research Institute, RRID:CVCL_C436) | 600 | 570 | Successful | Fused organoids were discarded from spinner flasks | Organoids were not frozen |
| 29 | Researcher 3 | Tubulin-GFP tagged hiPSCs (Coriell, Cat# AICS-0012) | 370 | 300 | Successful | Neurospheres lost during neural induction | Organoids were not frozen |
| 30 | Researcher 3 | Tubulin-GFP tagged hiPSCs (Coriell, Cat# AICS-0012) | 200 | 180 | Successful | Fused organoids were discarded | Organoids were not frozen |
| 31 | Researcher 3 | Tubulin-RFP tagged hiPSCs (Coriell Cat# AICS-0031-035) | 300 | 270 | Successful | Fused organoids were discarded | Organoids were not frozen |
| 32 | Researcher 4 | Tubulin-RFP tagged hiPSCs (Coriell Cat# AICS-0031-035) | 185 | 175 | Successful | Neurospheres lost during transfer | Organoids were not frozen |
| 33 | Researcher 4 | IMR90 (WiCell Research Institute, RRID:CVCL_C436) | 400 | 355 | Successful | Neurospheres lost during transfer | Organoids were not frozen |
| 34 | Researcher 4 | IMR90 (WiCell Research Institute, RRID:CVCL_C436) | 385 | 320 | Successful | Neurospheres lost during transfer | Organoids were not frozen |
| 35 | Researcher 4 | IMR90 (WiCell Research Institute, RRID:CVCL_C436) | 380 | 30 | Failed | Organoids were collapsed due to unknown reasons | Organoids were not frozen |
| 36 | Researcher 5 | IMR90 (WiCell Research Institute, RRID:CVCL_C436) | 380 | 300 | Successful | Neurospheres lost during transfer | Organoids were not frozen |
| 37 | Researcher 5 | IMR90 (WiCell Research Institute, RRID:CVCL_C436) | 300 | 270 | Successful | Neurospheres lost during transfer | Organoids were not frozen |
